# Ccr4-Not ubiquitin ligase signaling regulates ribosomal protein homeostasis and inhibits 40S ribosomal autophagy

**DOI:** 10.1101/2023.08.28.555095

**Authors:** Daniel L. Johnson, Ravinder Kumar, David Kakhniashvili, Lawrence M. Pfeffer, R. Nicholas Laribee

## Abstract

The Ccr4-Not complex containing the Not4 ubiquitin ligase regulates gene transcription and mRNA decay, yet it also has poorly defined roles in translation, proteostasis, and endolysosomal-dependent nutrient signaling. To define how Ccr4-Not mediated ubiquitin signaling regulates these additional processes, we performed quantitative proteomics in the yeast *Saccharomyces cerevisiae* lacking the Not4 ubiquitin ligase, and also in cells overexpressing either wild-type or functionally inactive ligase. Herein, we provide evidence that both increased and decreased Ccr4-Not ubiquitin signaling disrupts ribosomal protein (RP) homeostasis independently of reduced RP mRNA changes or reductions in known Not4 ribosomal substrates. Surprisingly, we also find that both Not4-mediated ubiquitin signaling, and the Ccr4 subunit, actively inhibit 40S ribosomal autophagy. This 40S autophagy is independent of canonical Atg7-dependent macroautophagy, thus indicating microautophagy activation is responsible. Furthermore, the Not4 ligase genetically interacts with endolysosomal pathway effectors to control both RP expression and 40S autophagy efficiency. Overall, we demonstrate that balanced Ccr4-Not ligase activity maintains RP homeostasis, and that Ccr4-Not ubiquitin signaling interacts with the endolysosomal pathway to both regulate RP expression and inhibit 40S ribosomal autophagy.

## INTRODUCTION

Cells dynamically respond to nutrient availability and stress through mechanisms that require communication between metabolic-responsive organelles, including the endolysosomal compartment and mitochondria, and the gene expression machinery (1,2). This communication allows cells to coordinate their growth and proliferation with favorable nutrient environments, while it has an equally important role in limiting anabolic and proliferative activity under unfavorable environments. Accumulating evidence indicates that dysregulating these processes causes many complex diseases and developmental syndromes, including cancer, neurodevelopmental disorders, aging, and metabolic disease (2). Deciphering these interrelationships remains an unrealized goal in molecular medicine as they are inherently complex with considerable functional redundancy.

The Ccr4-Not complex is a large multisubunit complex that is conserved throughout evolution and has critical roles at all stages of the gene expression process. Ccr4-Not originally was identified in budding yeast where it consists of the Ccr4, Caf1, Caf40, Caf130, and the Not1-5 subunits (3). The complex’s best understood role is regulation of cytoplasmic mRNA decay by the Ccr4 deadenylase subunit, and Ccr4-Not mediated mRNA degradation is highly conserved in metazoans (3,4). Genetic and biochemical studies also have shown that Ccr4-Not regulates RNA polymerase II (Pol II)-dependent transcription initiation, Pol II elongation, and mRNA nuclear export (3,5,6). Recently, Ccr4-Not was shown to regulate RNA Polymerase I (Pol I) transcription initiation and elongation of ribosomal RNA genes downstream of the target of rapamycin complex 1 (TORC1) signaling pathway that controls cell anabolism and proliferation (7). Although Ccr4-Not has important transcriptional activities, its effect on Pol I and Pol II transcription are less defined compared to its role in mRNA degradation.

While Ccr4 is the best understood enzymatic activity in the complex, Not4 is a highly conserved ubiquitin ligase whose *in vivo* functions remain poorly defined. Not4 has an N-terminal RING domain that interacts with the Ubc4 and Ubc5 E2 enzymes to mediate direct substrate ubiquitination (8). Among the few known Not4 substrates are the histone demethylase Jhd2 and the Cyclin C subunit of the Mediator complex, and their Not4-dependent ubiquitination stimulates proteasome-mediated degradation (9,10). Immediately C-terminal to the Not4 RING domain is a highly conserved RNA recognition motif (RRM) and C3H1 domain that are referred to collectively as the RRM-C domain (11). This domain has homology with similar domains found among other RNA binding proteins and, while Not4 cross-links to RNA *in vivo*, whether it binds RNA directly through the RRM-C domain is not clear. While Not4 RING mutants alone do not completely phenocopy cells lacking Not4 (8), combining both RING and RRM-C mutants simultaneously does phenocopy Not4 null cells (11). This combination of RING and RRM-C domains makes Not4 a completely unique ubiquitin ligase within the eukaryotic proteome (12). Besides direct substrate ubiquitination, Not4 also can function as an adaptor to mediate substrate ubiquitylation by additional E3 ubiquitin ligases, including the HECT-domain ligase Rsp5/NEDD4 (13). The capacity to recruit additional ubiquitin ligases such as Rsp5 may expand Ccr4-Not’s ubiquitin signaling activity into areas previously not associated with Ccr4-Not. For example, while Rsp5 regulates transcription and mRNA export (14–16), it also controls protein sorting and degradation through the endolysosomal compartment and it regulates autophagy (17–22).

Additional Not4 studies have revealed that it controls translation (23–25), proteostasis (26–28), and endolysosomal-dependent nutrient signaling (29), but its role in these processes remains incompletely understood. Specifically, Not4 monoubiquitinates both the ribosomal protein Rps7a and the Egd1 and Egd2 (Egd1/2) subunits that make up the nascent polypeptide associated complex (NAC) involved in co-translational quality control (23,25). This modification alters their function without signaling for their degradation. Loss of the Not subunits in Ccr4-Not, including Not4, reduces polysome formation and alters the ratios of the 40S, 60S, and 80S ribosomes relative to wild-type cells (23). Ccr4-Not physically interacts with ribosomes to facilitate co-translational assembly and translational repression (24,30,31), while it also monitors optimal codon usage during translation elongation (32), which involves the Not5 subunit and Not4-dependent Rps7a ubiquitination. Beyond translation, Not4 also maintains global proteostasis by ubiquitinating the 19S proteasome subunit Rpt5 (26,27). In the absence of Rpt5 ubiquitination, proteasomes assemble incorrectly, and they have deregulated 20S catalytic activity and defective 19S-dependent substrate deubiquitination (11,26–28). During extreme nutrient stress, these aberrant proteasomes also become sensitive to vacuole degradation through a specialized autophagy process termed proteaphagy (28). Cells lacking Ccr4 also deregulate autophagy gene transcripts to aberrantly activate autophagy (hereafter macroautophagy) in nutrient replete conditions (33). Macroautophagy activation results in the formation of a double membrane autophagosome that envelops cargo and then fuses with the vacuole to degrade the cargo and recycle the nutrients (34). Macroautophagy regulates protein quality control so Ccr4-Not, and specifically Not4, may contribute to nutrient stress responses and protein quality control through macroautophagy regulation. Ccr4-Not also activates nutrient signaling through the TORC1 pathway by promoting the function of the vacuole V-ATPase H^+^ pump that both acidifies the vacuole and activates TORC1 and PKA signaling in response to nutrients (29,35–37). Ccr4-Not mutants exhibit synthetic sick or lethal genetic interactions with endolysosomal pathway effectors, including V-ATPase mutants and vacuolar protein sorting (Vps factors) (8,38–40). Thus, Ccr4-Not is functionally connected to networks controlling translation, proteostasis, and endolysosomal-dependent autophagy and nutrient signaling. Yet how Ccr4-Not dependent ubiquitin signaling contributes to these processes remains incompletely understood.

The role individual Ccr4-Not subunits have on the transcriptome has been described previously (29,41–43), yet the effect that Not4 ligase signaling has on the proteome remains poorly understood. During the course of the study described herein, a report from Allen *et al*. used SILAC labeling coupled with mass spectrometry analysis of the differentially regulated proteome between WT and *not4Δ* cells (31). However, to our knowledge no analysis of the proteome has been performed in *not4Δ* reconstituted with either WT or inactive Not4, which would prevent changes to Ccr4-Not structure caused by the absence of Not4 that indirectly affect its non-ubiquitin signaling activities. We have addressed this deficiency by performing tandem mass tag (TMT) labeling and quantitative mass spectrometry analyses of the Not4 regulated proteome using a previously described vector system overexpressing either WT Not4 or a full-length Not4 mutant that phenocopies Not4 null cells. We find that both loss of Not4 ligase activity, or its overexpression, downregulates ribosomal proteins (RPs) and ribosomal biogenesis (Ribi) factors without reducing expression of known Not4 ribosomal substrates. This RP repression is independent of changes to RP mRNA expression, and it has functional consequences for both protein synthesis and maintaining proteostasis. Unexpectedly, we found that Ccr4-Not inhibits 40S ribosome autophagy, which is a specialized autophagy pathway whose selective regulators are unknown, in a Not4 ligase-dependent manner. Additionally, we provide genetic and functional evidence that this RP repression and 40S autophagy in Ccr4-Not ligase deficient cells is connected to factors that regulate the endolysosomal pathway.

## MATERIALS AND METHODS

### Yeast strains and culture conditions

All strains in this study are derived from the BY4741 genetic background and are listed in Table S1. For all experiments, cells were cultured in an incubated shaker at 30°C and harvested between OD_600_= 0.8-1.6. Cells were cultured in nutrient rich YPD (1% yeast extract/ 2% peptone/ 2% dextrose). To select for plasmid transformants, cells were cultured in synthetic complete (SC) media containing 0.2% yeast nitrogen base without amino acids, 0.5% ammonium sulfate, 2% dextrose, and 0.19% amino acid dropout mix with amino acids or uracil added back except the specific one needed for maintaining plasmid selection. The genetic manipulations used to either delete target genes, or to engineer genomically integrated in-frame epitope tags, were performed as described (44). All yeast plasmids used in this study are listed in Table S2.

### Chemical reagents

Most chemicals, including rapamycin, were purchased either through Fisher or Millipore Sigma. Yeast media was purchased from US Biologicals and 5-Fluoroorotic acid (5-FOA) was purchased from Research Products International.

### RNA extraction and RT-qPCR analysis

Total RNA was prepared by hot acid phenol extraction from three independent biological replicates per experimental condition. The ImProm-II reverse transcription kit (Promega) was used to synthesize cDNA from 1 µg of total RNA using the included oligo dT primers in a volume of 20 µL per the manufacturer’s direction. After cDNA synthesis, the cDNA reaction was diluted with ultrapure dH_2_O to a final volume of 100 µL, and 1 µL of the diluted cDNA was used to perform gene-specific qPCR using SYBR Green reagent and an ABI StepOne Plus thermal cycler. The gene specific signal was normalized to the *SPT15* internal reference control gene using the formula 2^(Ct*SPT15*-Ct*Target*)^ as previously described. The mean and standard deviation for each gene were calculated and are plotted in Figure S4 and Figure S7. These means then were Log_2_ converted and used to generate the heat map results for Figures 2 and Figure 5. Primer sequences used in this study are in Supplemental File 1. All statistical analyses and graphs were generated using GraphPad Prism 10.

### Antibodies and immunoblotting reagents

All immunoblots are representative of at least three independent experiments. The α-HA (clone F-7, catalog number sc7392), α-GFP (clone B-2, catalog number sc9996), and α-ubiquitin (clone P4D1, catalog number sc8017) antibodies were purchased from Santa Cruz Biotechnology. The α-FLAG (clone M2, catalog number F1804) and α-glucose-6-phosphate dehydrogenase (G6PDH, catalog number A9521) were purchased from Millipore Sigma. The α-puromycin antibody (clone PMY-2A4) was purchased from the Developmental Hybridoma Studies bank. Mouse and rabbit HRP-conjugated light chain specific secondary antibodies were purchased from Jackson ImmunoResearch Laboratories (catalog numbers 115-035-174 and 211-032-171). Immunoblots were performed using MilliporeSigma Immobilon^TM^ Western Chemiluminescent HRP Substrate and images were quantified using ImageJ.

### Whole cell extract preparation

Whole cell extracts (WCEs) were prepared by bead beating in extraction buffer (150 mM NaCl, 10 mM Tris, pH 8.0, 0.1% NP-40, 10% glycerol) and containing protease inhibitors (2 µg/mL each of aprotinin, pepstatin, and leupeptin), phosphatase inhibitor cocktail set II (Millipore Sigma, 1:100 dilution) and 1 mM DTT. Cell lysates were clarified for 15′ at 15,000 rpm @ 4°C, supernatants then were transferred to a new tube, and the protein concentration determined by Bradford assay.

### TUBE ubiquitin linkage analysis

WT and *not4Δ* Rps9a-GFP expressing cells were cultured to log phase in YPD media, harvested, and WCEs then prepared by bead beating in extraction buffer (150 mM NaCl, 10 mM Tris, pH 8.0, 0.1% NP-40, 10% glycerol, and 60 mM 2-chloroacetamide) containing protease inhibitors (2 µg/mL each of aprotinin, pepstatin, and leupeptin), and phosphatase inhibitor cocktail set II (Millipore Sigma, 1:100 dilution). For each sample, 600 µg of WCE was mixed with 2 µg of α-FLAG antibody in a total volume of 400 µL WCE buffer that had been mock treated or treated with 3 µg of K48ub or K63ub linkage specific Tandem Ubiquitin Binding Entities (TUBEs) purchased from LifeSensors. Samples were rotated at 4°C for a minimum of two hours before adding Protein A agarose beads and rotating at 4°C for an additional one hour to capture the immune complexes. Samples were washed three times with buffer and then resolved by 10% SDS-PAGE and analyzed by α-GFP immunoblotting.

### Tandem mass tag labeling and quantitative proteomic analysis

WCEs were prepared in 300 mM NaCl lysis buffer as described above. Samples (50ul, 50ug protein) then were reduced with 5 mM DTT for 45 min at 50°C, alkylated with 25 mM iodoacetamide for 20′ at room temperature in the dark, and then incubated with 20 mM DTT for an additional 15′ at room temperature. Samples then were precipitated with 5 volumes of cold acetone (at -20°C overnight), and proteins were pelleted at 16,000xg at 4°C for 10′. Protein pellets were washed with 100 ul of cold (-20°C) 90% acetone, air dried for four minutes, and re-dissolved in 100 µL of digestion buffer (100 mM HEPES, pH 8.3) containing 0.5 µg of Lys-C enzyme (Waco, Fuji, 125-05061). The protein mix then was digested for 2hrs at 37°C with shaking; adding 1ug trypsin (Thermo Fisher, 900057), the digestion was continued overnight.

For TMTpro-16plex labeling a set of 16 samples each containing 50 µg of peptides in 100 µL of digestion buffer (100 mm HEPES pH 8.3) was labeled using commercial TMTpro-16plex Mass Tag Labeling reagent kit (A44521, Thermo Fisher) according to the manufacturer’s protocol. The labeled set of 16 samples was combined, vacuum dried, and reconstituted in 0.1% TFA at 0.3 µg/µL for further fractionation. 300 µL (90 µg) of reconstituted mixture of labeled peptides was fractionated using Pierce High pH Reversed-Phase Peptide Fractionation kit (84868, Thermo Fisher) according to the manufacturer’s protocol for TMT-labeled peptides, with eight step fractions (consecutively eluted with 10.0, 12.5, 15.0, 17.5, 20.0, 22.5, 25.0, and 50.0% acetonitrile) collected. The peptide fractions were vacuum dried, dissolved in 65 µL of loading buffer (3% ACN with 0.1% TFA), and 5 µL (0.87 µg, assuming elution of equal amounts of peptides per fraction) aliquots were analyzed by LC-MS for peptide/protein identification and quantification.

Acquisition of raw MS data was performed on an Orbitrap Fusion Lumos mass spectrometer (Thermo Fisher) operating in line with Ultimate 3000RSLCnano UHPLS system (Thermo Fisher) using MS2 method for TMTpro-16plex labeled samples with 160 min LC gradient. The peptides were trapped on an Acclaim PepMap 100 nanoViper column (75 µm x 20mm, Thermo Fisher) at 5 µl/min flow rate and washed with loading buffer for 5min. The trapped peptides were separated on an Acclaim PepMap RSLC nanoViper column (75µm x 500mm, C-18, 2µm, 100Å, Thermo Fisher) at 300 nL/min flow rate and 40 °C column temperature using water and acetonitrile with 0.1% formic acid as solvents A and B, respectively. The following multi-point linear gradient was applied: 3% B at 0-4min, 5% B at 5′, 23% B at 110′, 30% B at 120′, 90%B at 123-133′, 3% B at 136-160′. MS2 acquisition method was used with three second cycles with the following MS scan parameters. Full MS scans were performed in the Orbitrap analyzer at 120,000 (FWHM, at m/z=200) resolving power to determine the accurate masses (m/z) of peptides. Following MS1 full scans, data dependent MS2 scans were performed on precursor ions with peptide isotopic pattern, charge state 2-6, and intensity of at least 25000. Peptide ions were isolated in the quadrupole with 0.7 m/z window, fragmented (HCD, 38% NCE), and the masses/intensities of fragment and reporter ions were determined with an Orbitrap analyzer at 50,000 (FWHM, at m/z=200) resolving power and dynamic exclusion applied for 45 seconds.

Post-acquisition analysis of raw MS data was performed within a mass informatics platform Proteome Discoverer 2.4 (Thermo Fisher) using Sequest HT search algorithm and yeast protein database (SwissProt, Saccharomyces cerevisiae S288c, TaxID 559292, v.2017-10-25, 6727 entries). The reversed target database was used as decoy database. The raw MS data acquired for the set of 8 fractions (derived from the same mixture of labeled samples) were treated as ‘Fractions’ for post-acquisition analysis. Full tryptic peptides were searched and two mis-cleavages were allowed. The searched fixed modifications included: carbamidomethylation of Cys residues and TMTpro modification of Lys residues and any peptide N-terminus. The variable modifications included oxidation of Met residues, acetylation, and Met-loss of the protein N-terminus. The precursor, fragment, and reporter ion mass tolerances were set to 10 ppm, 0.02 Da, and 20 ppm, respectively. The raw data were filtered for the precursor ions with S/N of at least 1.5. The PSMs were filtered for further analysis using a delta Cn threshold of 0.05. The q-values were calculated at PSM level (Percolator) and then, at the peptide level (Qvality algorithm), to control for the false discovery rate (FDR). The FDR threshold of 0.01 was used to validate and filter the PSMs and the corresponding peptide sequences. The validated/filtered peptides were used for the identification of the candidate precursor proteins. Each candidate precursor protein was scored by summing the PEP values of the assigned peptides. The sum-PEP protein scores were used to calculate the experimental q-values at the protein level. The candidate proteins were further validated using 0.01 (strict) and 0.05 (relaxed) FDR thresholds. A set of candidate proteins was accepted as an identified protein group if none of the assigned peptides was unique to any protein, but at least one of those peptides was common and unique to that group of proteins. Strict parsimony principle was applied to the protein groups. The reporter ions quantification values (based on S/N ratio) were corrected according to the product data sheet provided with the used TMTpro-16plex Label reagent set. The co-isolation and average S/N ratio thresholds were set to 50% and 10, respectively. Unique and razor peptides were used for protein quantification, and protein groups were considered for peptide uniqueness.

### Proteome data analysis

The data generated by the mass spectrometry analysis was transferred to the UTHSC Molecule Bioinformatics Core using SFTP. These data then were normalized using normalizeCylclicLoess function from R/Bioconductor-package limma (45). The normalized data matrix was loaded into R, and statistics were gathered and used to determine differential expression. The mean, variance, standard deviation, and standard error of means were calculated for each protein across the different conditions. Pearson’s correlation coefficient was graphed in order to identify sample outliers in each condition. At this time no outliers were determined. A principal component analysis was performed to determine different clusters in the samples, and fold change was then calculated for all proteins. The significance for each protein was assessed by using one-way ANOVA, and the p values were adjusted for multiplicity using the Benjamini Hochberg method (46). Only proteins with an adjusted p value < 0.05 were considered differentially expressed (Supplemental File 2). These proteins are used to create a heatmap with unsupervised hierarchical clustering. The protein list then was loaded into the STRING database for protein-protein interaction mapping and to test for enriched cellular component GO categories (47).

## RESULTS

### Both increased Not4 dosage, and loss of its ligase activity, inhibit RP and Ribi protein expression

Although Ccr4-Not regulation of the transcriptome has been defined in-depth, the impact that Ccr4-Not ligase activity has on the proteome remains incompletely defined. Furthermore, most Ccr4-Not ligase studies have used cells either lacking Not4 expression entirely (*not4Δ*) (31), express Not4 partial deletion constructs (13), or have reconstituted *not4Δ* with Not4 RING mutants that do not completely phenocopy *not4Δ* (8). Accurate assessment of Not4 function *in vivo* necessitates using a full-length Not4 ligase that incorporates into Ccr4-Not yet phenocopies *not4Δ* to avoid any possible indirect effects on Ccr4-Not activity that are independent of its ubiquitin ligase role. To address these concerns, we transformed wild-type (WT) and *not4Δ* with control vector, vector overexpressing FLAG-tagged WT Not4 (*NOT4*), or vector expressing a Not4 mutant (*NOT4RR*) that contains mutations in both the RING and RRM-C domains. The Not4RR incorporates into Ccr4-Not yet it phenocopies a *not4Δ* . The plasmid expressed *NOT4* restored all *not4Δ* phenotypes, including sensitivity to heat stress, while *NOT4RR* failed to restore any *not4Δ* phenotype tested (Figure 1A and see below). Importantly, *NOT4* and *NOT4RR* transcripts are expressed at significantly higher levels relative to *NOT4* expressed from its endogenous chromosomal locus (Figure 1B). We reasoned this Not4 overexpression system could be used to identify proteins whose expression is sensitive to Not4 dosage and/or its ligase activity, which could facilitate identification of possible Not4 substrates and novel Ccr4-Not ligase regulated pathways.

**Figure 1.**
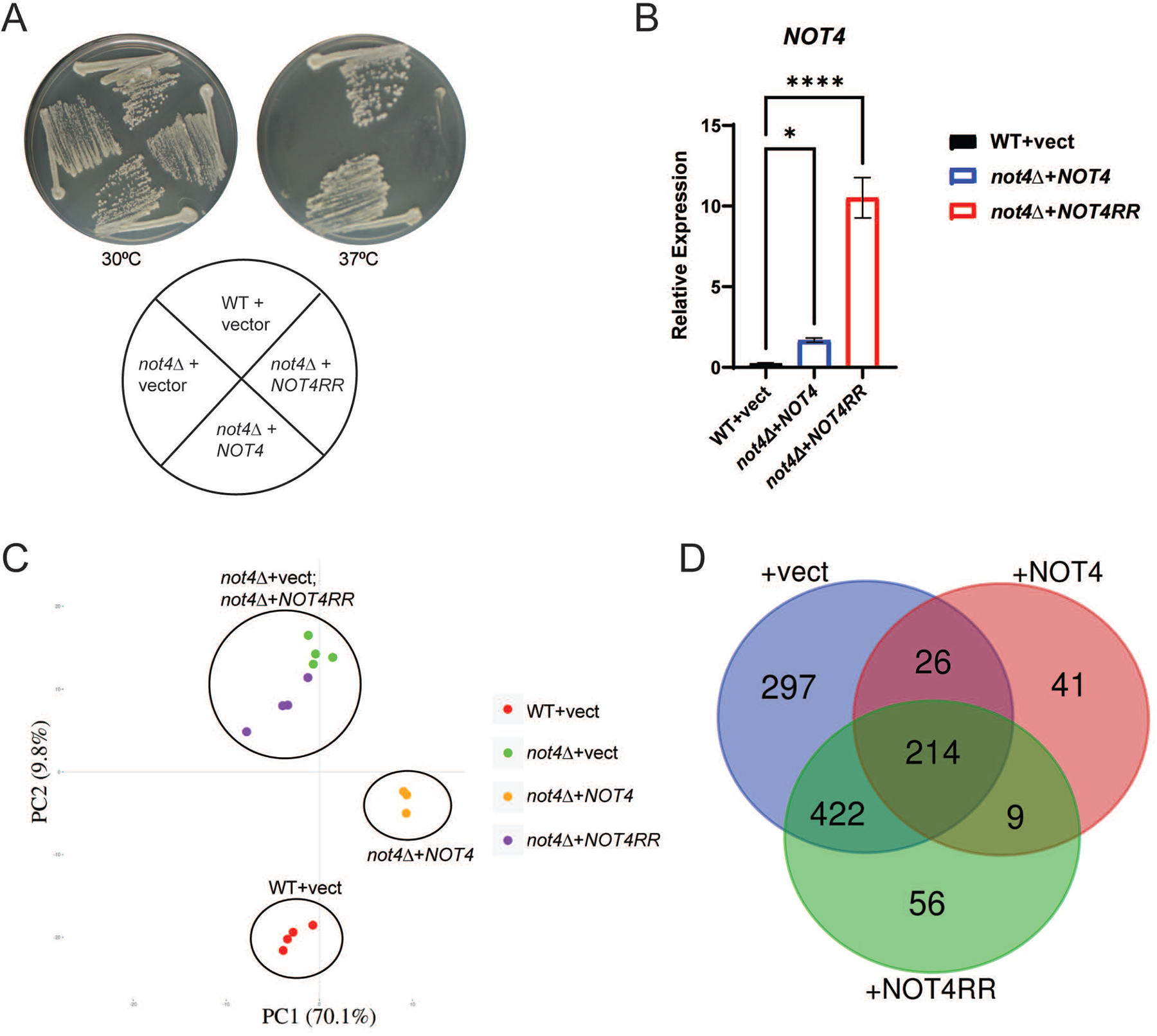
Not4 ligase dosage and activity differentially regulate the cellular proteome. **A.** Wild-type (WT) cells transformed with control vector, and *not4Δ* cells expressing either control, *NOT4*, or *NOT4RR* vector, were streaked to duplicate plates and incubated at control temperature (30°C) or heat shock temperature (37°C) for three days before image capture. **B.** Reverse transcriptase coupled with quantitative PCR (RT-qPCR) analysis of *NOT4* transcript levels from the indicated strains. The *NOT4* specific signal was normalized to the expression of the internal control gene *SPT15* and significance was tested by one-way ANOVA. *\*-p< 0.05* or greater. **C.** Proteome principal component analysis (PCA) of the individual replicates for the indicated strains. Each experimental condition had four independent biological replicates except for *not4Δ* + *NOT4*, which had three independent biological replicates. **D.** The differentially expressed proteins (DEPs) were identified by using a stringent 1.5-fold change in expression and an FDR < 0.05. The resulting DEPs then were subjected to Venn analysis. Total number of DEPs in each condition relative to the WT + vector control is 959 (*not4Δ* + vector), 290 (*not4Δ* + *NOT4*), and 701 (*not4Δ* + *NOT4RR*).

To identify differentially expressed proteins, whole-cell extracts (WCEs) from at least three (*not4Δ* + *NOT4*) independent biological replicates from Figure 1A were prepared and subjected to LC-MS-MS analysis using Reporter Ions Quantification approach based on Tandem Mass Tag (TMTpro)-labeling. Quantifiable results for over 3,300 proteins in each condition was achieved, and principal component analysis (PCA) revealed high reproducibility between the independent replicates as evidenced by their clustering together for each condition (Figure 1C). Intriguingly, while *not4Δ* + *NOT4* restored *not4Δ* phenotypes (Figure 1A), the *not4Δ* + *NOT4* clusters distinctly from the WT + vector control, thus indicating that Not4 overexpression results in differences in their proteomes (Figure 1C) . The *not4Δ* + vector and *NOT4RR* samples clustered similarly (Figure 1C), which is predicted since the Not4RR mutant phenocopies a *not4Δ* (Figure 1A) . To identify the differentially expressed proteins (DEPs) between the three experimental conditions, the data were analyzed using a stringent 1.5-fold change in expression with an FDR<0.05 relative to the WT + vector control. Using these criteria, 959 (*not4Δ* + vector), 290 (*not4Δ* + *NOT4*), and 701 (*not4Δ* + *NOT4RR*) DEPs were identified (Figure 1D). Venn analysis of these DEPs identified substantial overlap (214 proteins) between the vector, *NOT4*, and *NOT4RR* expressing cells, which indicates a population of proteins whose expression is highly sensitive to both Not4 dosage and its ligase activity (Figure 1D). Pairwise analyses also reveal exclusive overlap between the vector and *NOT4* (26 proteins), *NOT4* and *NOT4RR* (9 proteins), and vector and *NOT4RR* (422 proteins) (Figure 1D). Additionally, the *not4Δ* vector, *NOT4*, or *NOT4RR* cells exhibit condition-specific DEPs, with 297 DEPs specific to the *not4Δ* vector, while 41 and 56 DEPs were selective for *NOT4* or *NOT4RR*, respectively (Figure 1D). These data indicate that both Not4 dosage and its ligase activity have both overlapping and selective effects on the proteome.

The total DEPs for each experimental condition were subjected to cellular component Gene Ontology (GO) analysis through the STRING database to determine if any specific functional networks were overrepresented. For the *not4Δ* + vector, the GOs were dominated by categories related to the nucleolus, preribosome, and cytoplasmic ribosome (Figure 2A). When these DEPs were further sub-divided into up and down DEPs and analyzed separately, the down DEPs were specific for the nucleolar and ribosomal-related GOs while the up DEP GOs were enriched for proteasome regulatory particle, polarized growth, and cytoskeletal functions (Figure S1). Ribosomal-specific GOs also were overrepresented in *NOT4* expressing cells, although not to the same extent as in the *not4Δ* vector cells. Additionally, a substantial number of the *NOT4* GOs were related to mitochondrial function and metabolism (Figure 2B). Parsing these results further revealed that the GOs for the *NOT4* down DEPs all related to ribosomal function, while the up GOs were mitochondria-specific (Figure S2). These data indicate that increased Not4 dosage reduces ribosomal factors but enhances expression of mitochondrial proteins. Similar analysis of the aggregate *not4Δ + NOT4RR* DEPs revealed nucleolar, ribosomal, and cytoplasmic stress granule GOs were overrepresented (Figure 2C). Finer resolution analysis revealed the down DEPs to be specific for ribosomal GO categories, while no specific GO categories were overrepresented in the up DEPs (Figure S3). Collectively, these proteomic data indicate that both excess Not4, or loss of Not4 ligase activity, represses RPs and ribosomal biogenesis (Ribi) factors.

**Figure 2.**
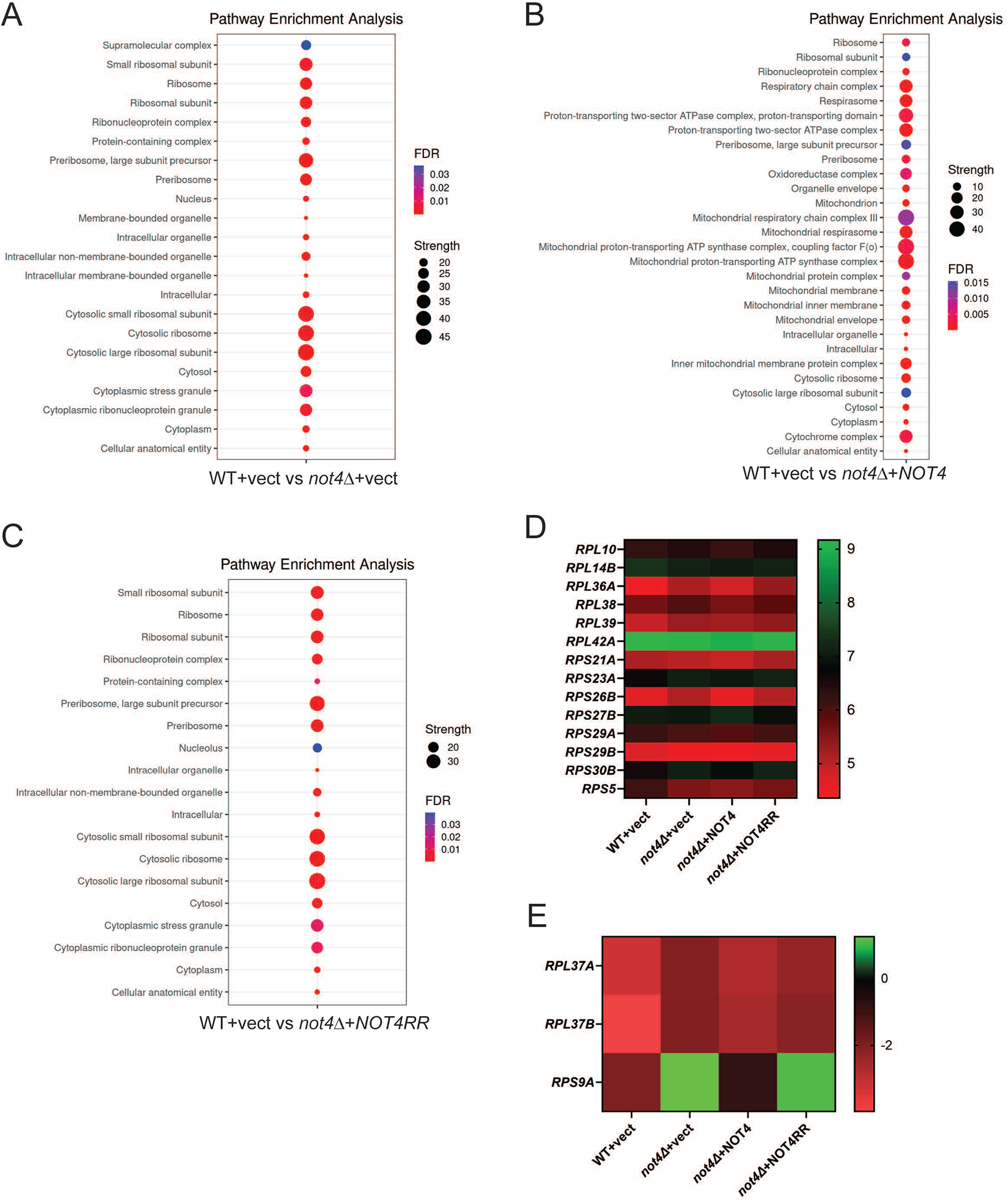
Ribosomal protein (RP) and ribosomal biogenesis (Ribi) factors are repressed by changes to Not4 activity or dosage. **A-C**. Dot blot analysis of the top cellular component gene ontology (GO) categories for the indicated experimental conditions. GO analysis was performed through the STRING database. Supplemental File 3 has the GO category and individual protein results used to generate these dot blots. **D** and **E**. RT-qPCR analysis of the indicated RP mRNAs was performed from three independent biological replicates normalized to the control gene *SPT15*. The average signal for all three replicates was Log_2_ converted and plotted in heat map format. Individual bar graphs for the expression of each RP mRNA with standard deviations plotted are presented in Figure S4.

Since RPs constitute the functional ribosome, we specifically examined whether the RP repression was due to inhibition of their corresponding RP mRNAs. RP genes are coordinately regulated, so we chose a diverse number of RPs from the proteomic dataset and performed RT-qPCR to quantify their mRNA levels. A minor, but statistically significant (*p< 0.05* by Student’s t-test), decrease in *RPS5* mRNA was found in the *not4Δ* vector, *NOT4*, and *NOT4RR* cells relative to the WT vector control (Figure 2D and Figure S4). However, the expression of the remaining RP mRNAs either was unaffected or upregulated in these cells relative to the WT vector control (Figure 2D-2E and Figure S4 for individual bar graph results). As such, these data indicate that RP inhibition in the *not4Δ* vector, *NOT4*, and *NOT4RR* cells cannot be explained by decreased expression of RP mRNAs. Not4 contributes to translational regulation in part by monoubiquitinating the RP Rps7a and the nascent polypeptide associated complex (NAC) factors Egd1 and Egd2 (Egd1/2) to promote co-translational quality control (23,25). The proteome data revealed that Rps7a and Egd1/2 expression was unaffected in all three experimental conditions (less than 1.5-fold change in abundance, see Figure S5). Therefore, the global RP inhibition in cells either completely lacking Not4, expressing an inactive Not4, or overexpressing Not4, is not due to repression of known Not4 translational substrates.

### Loss of Not4 ligase activity impairs protein synthesis and induces 40S RP degradation

Ccr4-Not mutants, including *not4Δ*, reduce polysome formation and alter the relative ratios of 40S and 60S ribosomes (23). To test if the RP repression detected was associated with effects on protein synthesis, the WT control, and the *not4Δ* vector, *NOT4*, and *NOT4RR* expressing cells, were metabolically labeled with puromycin and samples were taken at 0’, 30’, and 60’ timepoints. Puromycin incorporates into nascent protein chains blocks protein synthesis and can be detected by α-puromycin immunoblot (IB), which indicates active protein synthesis (48). A time-dependent increase in puromycin incorporation was readily detectable in the WT vector control cells, while puromycin incorporation was reduced dramatically in both the *not4Δ* vector and *NOT4RR* samples (Figure 3A). Although *NOT4* overexpression inhibits a large network of RPs and Ribi factors (Figure 2B and Figure S6), *NOT4* restored protein synthesis back to the WT state (Figure 3A). Therefore, the ribosomal inhibition in *NOT4* cells is insufficient to impair protein synthesis, while protein synthesis in *not4Δ* and *NOT4RR* expressing cells is profoundly inhibited.

**Figure 3.**
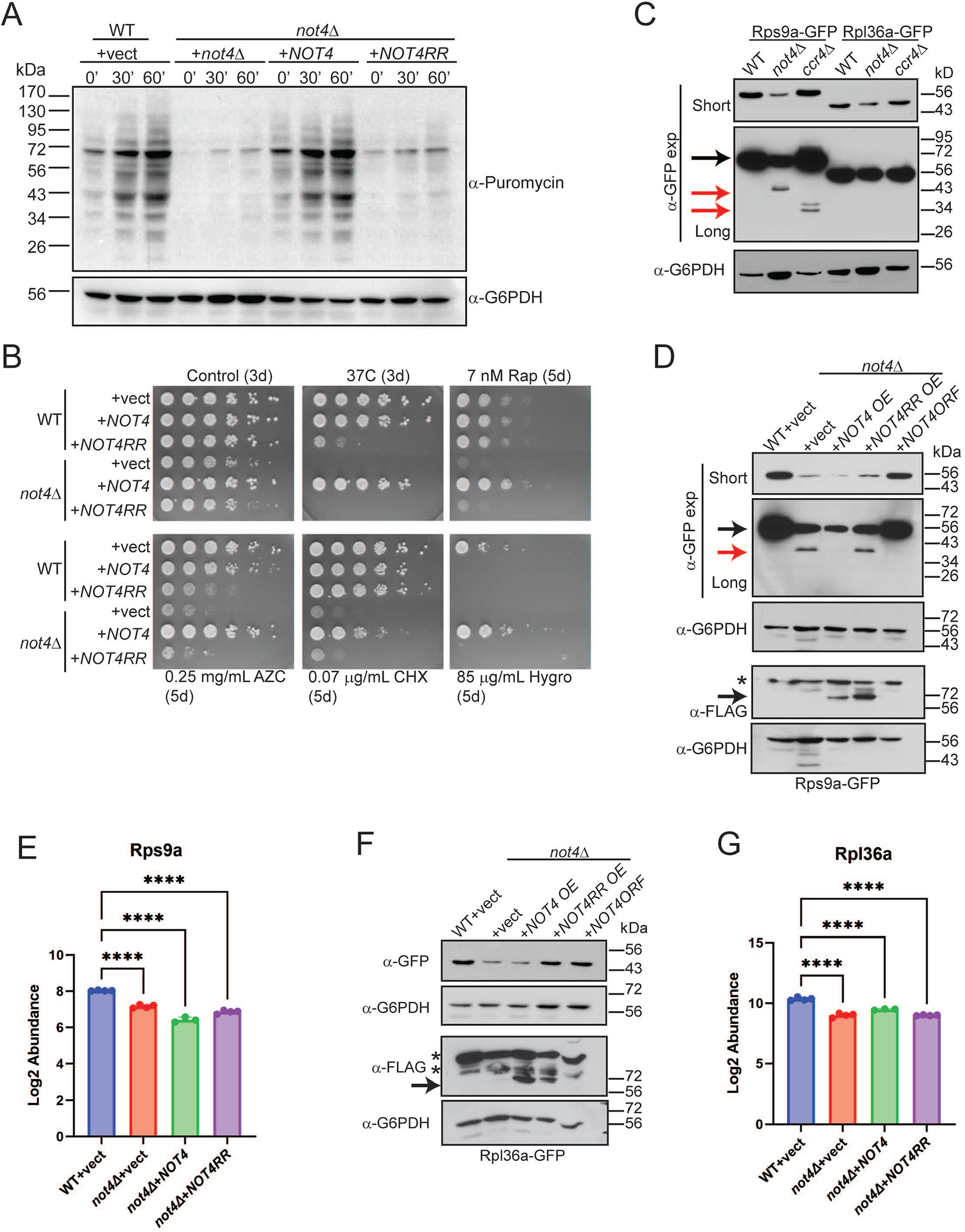
Both Not4 ligase activity and dosage repress RP expression to impact ribosomal activity *in vivo*. **A.** The indicated cells were grown to log phase and labelled with 10 µM puromycin before taking samples at the indicated timepoints. Whole cell extracts (WCEs, 30 µg) then were analyzed by α-puromycin immunblot (IB). The α-G6PDH IB serves as a loading control. **B.** WT and *not4Δ* transformed with the indicated expression vectors were cultured overnight at 30°C. Equal numbers of cells then were 5-fold serially diluted and spotted onto the indicated plates. The number of days the plates were incubated at either 30°C or 37°C before pictures were taken are indicated in parentheses. **C.** WCEs (30 µg) from the indicated strains in the Rps9a-GFP or Rpl36-GFP reporter backgrounds were analyzed by α-GFP and α-G6PDH IB. **D.** Duplicate IBs of WCEs (30 µg) from the indicated strains were probed with α-GFP to detect Rps9-GFP and α-FLAG to detect overexpressed Not4 or Not4RR. The α-G6PDH IBs serve as the loading control for each. Short and long exposures for the α-GFP IB are presented. Black arrow indicates full-length Rps9a-GFP and red arrow indicates partially degraded Rps9-GFP. The * in the α-FLAG IB indicates a non-specific band. **E.** The mean and SD of the peptide abundance signal from the proteomic data for Rps9a of the individual replicates. *\*\*\*\*-p<0.001* by one-way ANOVA. The Log_2_ fold-change (FC) for the individual *not4Δ* samples relative to the WT + vector are as follows: +vector (-0.87); +*NOT4* (-1.6); +*NOT4RR* (-1.2). **F.** Same as in **D** except Rpl36a-GFP is analyzed. **G.** The mean and SD of the peptide abundance signal from the proteomic data for Rpl36a of the individual replicates. *\*\*\*\*-p<0.001* by one-way ANOVA. The Log_2_ fold-change FC for the individual *not4Δ* samples relative to the WT + vector are as follows: +vector (-0.87); +*NOT4* (-1.6); +*NOT4RR* (-1.2).

The impact Not4 dosage and ligase activity have *in vivo* was probed further by analyzing their sensitivity to proteostatic stress (37°C heat stress and azetidine-2-carboxylic acid (AZC) which causes protein folding stress), nutrient stress (the TORC1 inhibitor rapamycin), or translational stress (cycloheximide and Hygromycin B) (11). For an additional comparison, WT cells also were transformed with the *NOT4* or *NOT4RR* expressing plasmids. As previously reported, *not4Δ* vector and *NOT4RR* cells exhibited marked sensitivity to the stress conditions tested, while *NOT4* rescued all of these phenotypes (Figure 3B). In WT cells, *NOT4RR* sensitized cells to proteostatic stress (heat stress and AZC), but it had no effect on nutrient stress or cycloheximide-induced translational stress (Figure 3B). These results indicate that Not4RR likely functions as a dominant negative in WT cells by competing with endogenous Not4 for Ccr4-Not incorporation, and they suggest that Ccr4-Not’s role in proteostasis is more sensitive to Not4 ligase activity compared to other stress responses. Intriguingly, while *NOT4* expression had no effect on the sensitivity of WT cells to most stressors, both *NOT4* and *NOT4RR* profoundly inhibited WT growth in the presence of Hygromycin B (Figure 3B). Hygromycin B stabilizes tRNA interactions with the ribosomal A-site to inhibit tRNA translocation (49), while cycloheximide binds the ribosomal 60S E-site to block ribosome elongation (50). These genetic results indicate that both increased and decreased Not4 ligase activity negatively affect translational control through mechanisms that may involve tRNA-ribosome interactions. Overall, these results indicate that Not4-dependent proteostasis regulation is highly sensitive to Not4 activity relative to other stress responses, and that Not4 dosage selectively sensitizes cells to distinct translational inhibitors.

To further delineate Not4-dependent effects on RP expression, we chose two representative RPs, the 40S subunit Rps9a and the 60S subunit Rpl36a, contained in the 214 overlapping DEPs for all three conditions (Figure 1D). An EGFP epitope tag was integrated in-frame at their genetic loci in WT, *not4Δ*, and *ccr4Δ* such that Rps9a and Rpl36a are expressed as in-frame C-terminal GFP fusions from their native promoters. Using these reporter strains, we found *not4Δ* reduces the expression of both RPs as predicted from the proteomic results, while *ccr4Δ* had no effect on either (Figure 3C, short α-GFP exposure). These data confirmed that RP inhibition in *not4Δ* is independent of Ccr4-dependent transcription or mRNA deadenylation. While Rps9-GFP is ∼ 56 kDa, longer exposure of the α-GFP IB revealed a discrete, faster migrating Rps9a-GFP band in *not4Δ* at ∼ 40 kDa and a doublet at ∼ 27-34 kDa in *ccr4Δ* (Figure 3C). These smaller Rps9a-GFP bands are absent from the WT, and smaller RP-specific bands are not present in the WT, *not4Δ*, or *ccr4Δ* Rpl36a-GFP expressing cells (Figure 3C). A recent report demonstrated that *ccr4Δ* upregulates basal macroautophagy in repressive (nutrient-rich) conditions (33). Autophagy flux is assessed in yeast by fusing GFP to candidate autophagy substrates and then monitoring release of free GFP (∼ 27 kDa) that occurs when autophagy cargoes are degraded in the vacuole since GFP resists vacuolar degradation (51). The faster migrating Rps9a-GFP specific bands indicate the possibility that in these autophagy repressive conditions, loss of either Not4 or Ccr4 increases 40S, but not 60S, ribosomal autophagy (referred to hereafter as ribophagy) (52).

The effect that Not4 ligase activity and dosage has on these specific RPs also was assessed directly by transforming the WT and *not4Δ* Rps9a-GFP and Rpl36a-GFP reporter strains with control, *NOT4*, or *NOT4RR* expressing plasmids. Separately, a different plasmid also was transformed into *not4Δ* that expresses *NOT4* from its endogenous promoter (*NOT4ORF*) so that *NOT4* expression most closely mirrors that of WT cells. Rps9a-GFP remained repressed in the *not4Δ* vector, *NOT4*, and *NOT4RR* cells (Figure 3D), which replicates the results from the proteomic analysis (Figure 3E), while the *NOT4ORF* restored Rps9a-GFP back to WT levels (Figure 3D). Furthermore, the partial Rps9a-GFP degradation product in the *not4Δ* vector and *NOT4RR* samples was absent in both *NOT4* and *NOT4ORF* cells even though *NOT4* failed to restore Rps9a-GFP expression (Figure 3D). For both *not4Δ* vector and *NOT4*, Rpl36a-GFP was repressed compared to WT (Figure 3F), which is consistent with our proteome results (Figure 3G), while *NOT4ORF* restored Rpl36a-GFP levels back to the WT state (Figure 3F). Interestingly, *NOT4RR* also consistently restored Rpl36a-GFP back to near WT levels, which differs from our proteome results (Figure 3G). The reason for this difference currently is unclear, but it may be due to unexpected effects the GFP tag has on Rpl36a. Collectively, these data indicate that a narrow range of Not4 ligase activity is required for maintaining wild-type RP expression, and these RP effects encompass both 40S and 60S RPs. They also indicate that the partial Rps9a-GFP degradation in *not4Δ* specifically results from lack of Not4 ligase activity independently of the Not4-dependent Rps9a repression.

### Not4 ligase loss represses 40S RP expression independent of proteasome-mediated degradation but dependent on the endolysosomal pathway

Recent studies have determined that loss of Not4-mediated ribosomal Rps7a monoubiquitination causes defects in protein solubility and increases protein aggregate formation (23,24). To test if this possibility could explain our results, equal numbers of WT and *not4Δ* cells expressing Rps9a-GFP or Rpl36a-GFP were lysed by boiling in SDS-PAGE sample buffer and then analyzed by α-GFP IB. Even when samples were prepared under denaturing conditions, *not4Δ* still repressed both RPs (Figure 4A). Importantly, since overexpressing WT Not4 represses a large network of RP and Ribi factors (Figure 2 and Figure 3), their inhibition cannot be explained by protein insolubility due to aggregate formation.

**Figure 4.**
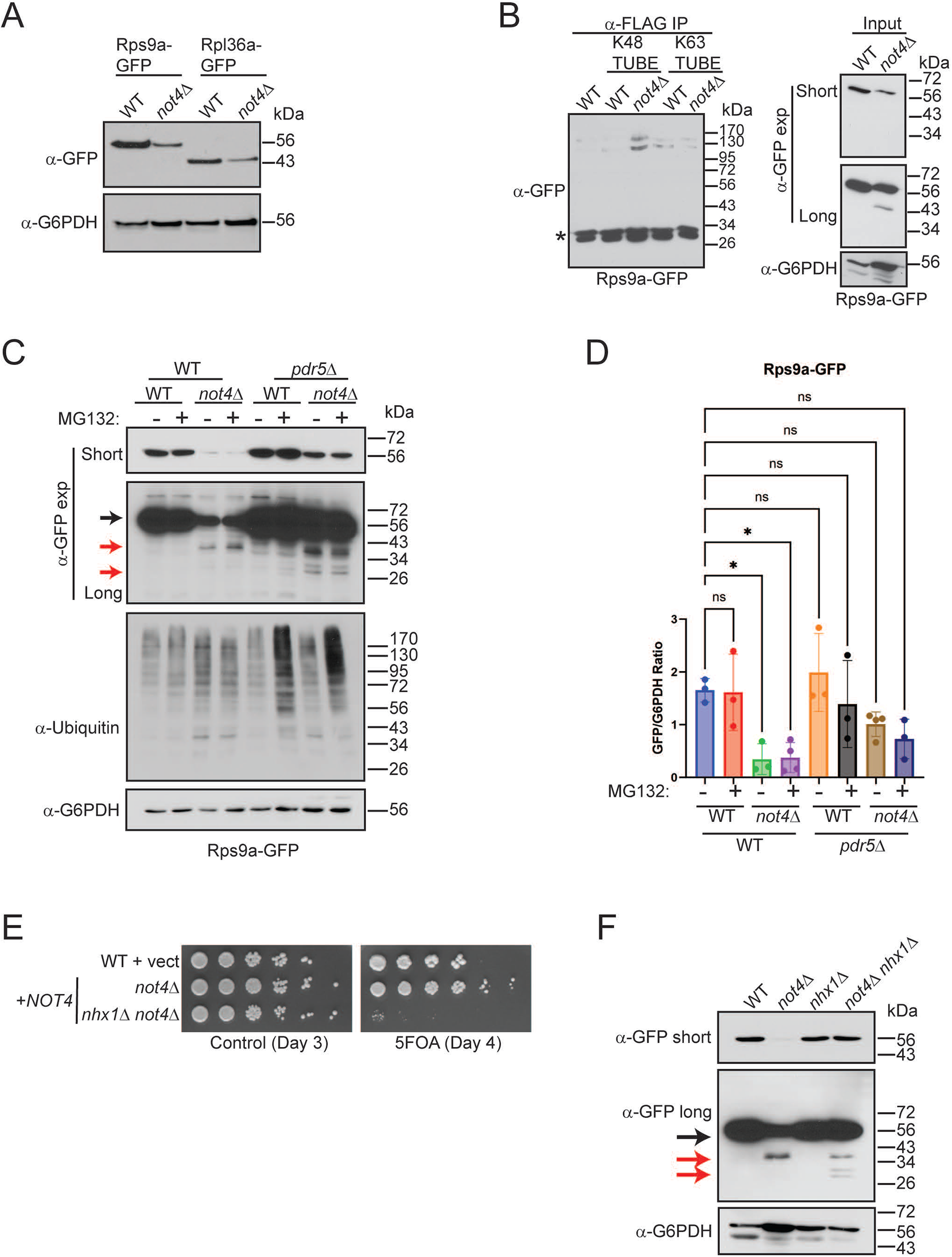
Not4 ligase loss causes Rps9a degradation independently of the proteasome but dependent on the endolysosomal pathway. **A.** Equal numbers of WT and *not4Δ* were lysed by boiling in SDS-PAGE loading buffer and then analyzed by α-GFP IB. **B.** All WCEs (600 µg) from the indicated strains were treated with α-FLAG antibody combined in the absence or presence of 3 µg of either K48 or K63-specific TUBES. After incubation with rotation at 4°C for two hours, protein A conjugated agarose beads were added and incubated for an additional one hour. The bead-immune complexes then were pelleted and washed extensively before analyzing by 10% SDS-PAGE and α-GFP IB. The inputs are 30 µg of the WCEs. Short and long α-GFP exposures are provided. The α-G6PDH IB serves as the loading control. *-non-specific band. **C.** The indicated strains were grown to log phase and then mock treated or treated with 10 µM MG132 for two hours to inhibit the proteasome. Short and long α-GFP IBs are provided. Black arrow indicates full-length Rps9a-GFP and red arrows indicate partial and complete degradation products. The α-G6PDH IB is the loading control. **D.** Quantitation of **C**. The short α-GFP IBs for a total of 3-4 individual biological replicates were quantified and expressed as the ratio of full-length GFP to G6PDH. *\*-p< 0.05* by one-way ANOVA. **E.** The WT and *not4Δ* strains were transformed with control or *NOT4* expression vector. Equal numbers of cells from overnight cultures then were 5-fold serially diluted and spotted to either plasmid-selective media or media containing 5-FOA and images were captured on the indicated days. **F.** Log phase WCEs (30 µg) from the indicated WT, single, and double mutants were IB with α-GFP. Short and long exposures are provided with the black arrow denoting full-length Rps9a-GFP and the red arrows indicating degradation products. The α-G6PDH IB is the loading control.

Not4 loss impairs proteosome assembly, deregulates its catalytic activity, and disrupts proteasome-dependent ubiquitin recycling which increases global protein ubiquitination (11,26–28). RPs produced in excess beyond that needed for ribosome biogenesis are flagged with ubiquitin and degraded through the ubiquitin-proteasome (UPS) system (53,54). Using Rps9a as a model RP, we assessed if Rps9a was ubiquitinated in *not4Δ* to promote RP inhibition through proteasome-mediated degradation. Initially, WCEs from WT and *not4Δ* were treated with FLAG-tagged K48 or K63 linkage-specific Tandem Ubiquitin Binding Entities (TUBEs) before performing α-FLAG pull-down assays and Rps9a detection by α-GFP IB (55). While only background levels of a high molecular weight (∼ 130-170 kDa) Rps9a-GFP specific signal was detected in the control and WT K48-TUBE specific IPs, the Rps9a-GFP signal increased specifically in the *not4Δ* K48-TUBE IP (Figure 4B). Separately, the K63-TUBE IPs pulled down K63-linked ubiquitinated Rps9a above background from WT, yet *not4Δ* did not alter its abundance substantially (Figure 4B). These data suggest that Not4 loss increases K48-linked Rps9a ubiquitination. Importantly, the high molecular weight Rps9a-GFP signal is specific since Rps9a-GFP signal was undetectable in the normal Rps9a-GFP size range of ∼ 56 kDa, thus indicating Rps9a likely is modified by multiple K48 and K63 specific ubiquitin chains (Figure 4B).

K48-linked ubiquitin targets proteins for proteasomal degradation, so we tested if the increased K48-linked ubiquitin on Rps9a in *not4Δ* flags it for proteasome-dependent RP degradation. WT and *not4Δ* were mock treated or treated for two hours with 10 µM MG132 to inhibit the proteasome. Since WT cells are resistant to proteasome inhibitors, we also engineered a *pdr5Δ* into both WT and *not4Δ*. Pdr5 is a plasma membrane efflux pump that exports both xenobiotics and H^+^ from cells (56), and its loss sensitizes cells to proteosome inhibitors (57). While MG132 treatment did not inhibit proteasome activity in WT or *not4Δ*, *pdr5Δ* loss sensitized both WT and *not4Δ* to proteasome inhibition as indicated by the MG132-dependent increase in global protein polyubiquitination (Figure 4C). Unexpectedly, we found that *not4Δ pdr5Δ* restored Rps9a expression back to near WT levels irrespective of proteasome inhibition (Figure 4C and Figure 4D). Longer α-GFP exposure also revealed that *not4Δ pdr5Δ* enhanced Rps9a-GFP degradation relative to *not4Δ* such that some detectable free GFP (∼ 27 kDa) accumulated, which was not increased further by proteasome inhibition (Figure 4C). These data indicate that *not4Δ* does not repress this 40S RP, or promote its incomplete degradation, through a proteasome-dependent mechanism. They also indicate that Pdr5 loss in *not4Δ* opposes the inhibitory effect on Rps9a expression while it facilitates Rps9a degradation.

Previously, we found that loss of either Not4 or Ccr4 inhibits TORC1 signaling by inhibiting vacuole V-ATPase activity and reducing vacuole acidity (29). The V-ATPase is essential for vacuole-dependent degradation of autophagy substrates, nutrient storage and recycling, and the downstream activation of RP and Ribi gene transcription and mRNA translation necessary for ribosomal biogenesis (58,59). Since Pdr5 can export H^+^, we speculated that Pdr5 may oppose vacuole V-ATPase activity indirectly by competing for a limiting pool of free cytoplasmic H^+^ (60). If so, then in Ccr4-Not deficient cells where V-ATPase function is partially compromised (29), Pdr5 loss may alleviate the inhibitory effect that Ccr4-Not disruption has on V-ATPase activity by reducing competition for H^+^. To independently test this possibility, *not4Δ* was combined with a *nhx1Δ* mutant lacking the vacuole Na^+^-K^+^/H^+^ exchanger that prevents vacuole H^+^ release and results in vacuole hyperacidification (61). While *not4Δ nhx1Δ* reconstituted with *NOT4* from a *URA3* expression plasmid had no overt growth defects, selection on 5-FOA media to force *NOT4* plasmid loss revealed the *not4Δ nhx1Δ* to be synthetically sick compared to *not4Δ* (Figure 4E). However, the *not4Δ nhx1Δ* fully restored Rps9a-GFP expression, and it increased conversion of the partial Rps9-GFP degradation product (∼ 40 kDa) to free GFP (∼ 27 kDa) (Figure 4F). These results suggest that Ccr4-Not disruption may repress RP expression and inhibit 40S RP degradation by disrupting V-ATPase dependent endolysosomal regulation. Furthermore, the RP repression in *not4Δ* cannot be explained as an indirect consequence of poor growth since the *not4Δ nhx1Δ* restores Rps9a expression yet it is more profoundly growth impaired (Figure 4E).

### 40S ribophagy activation in Ccr4-Not ligase-deficient cells is independent of TORC1 inhibition and the Atg7-dependent macroautophagy pathway

The above results indicate that the increased K48-linked RP ubiquitination in *not4Δ* does not signal for RP proteasomal degradation. Combined with the increased partial (in *not4Δ*) and complete (in *ccr4Δ*) Rps9a-GFP degradation shown in Figure 3C, we considered the alternative possibility that Ccr4-Not disruption represses RP levels by a macroautophagy-dependent mechanism. We initially tested if macroautophagy deregulation occurs in *not4Δ* as was reported for *ccr4Δ* by transforming WT, *ccr4Δ*, and *not4Δ* with a GFP-Atg8 expressing plasmid. GFP is cleaved from Atg8 when the autophagosome fuses with the vacuole, and since GFP resists vacuole degradation, free GFP accumulation indicates increased macroautophagy (51). Log phase cultures of two independent plasmid transformants from each background were analyzed by α-GFP IB and, consistent with macroautophagy inhibition in nutrient-rich environments, WT cells have no detectable free GFP (Figure 5A). As previously reported, *ccr4Δ* increased macroautophagy in nutrient-rich conditions as indicated by the presence of free GFP (33), while *not4Δ* also caused a similar level of macroautophagy activation (Figure 5A). The *ccr4Δ* deregulates autophagy by stabilizing autophagy (*ATG*) transcripts, so we tested if loss of Not4 ligase signaling activity also enhances *ATG* transcript levels. Expression of *ATG* mRNAs either were unaffected or elevated in *not4Δ* vector and *NOT4RR* cells, while their expression in *NOT4* was comparable to the WT control or only slightly upregulated (Figure 5B and Figure S7). Thus, both Ccr4-Not ubiquitin ligase and deadenylase activities repress expression of *ATG* transcripts in macroautophagy-inhibitory environments. However, it currently is unknown if Ccr4-Not ligase signaling represses these *ATG* transcripts by promoting their mRNA decay or by inhibiting their transcription.

**Figure 5.**
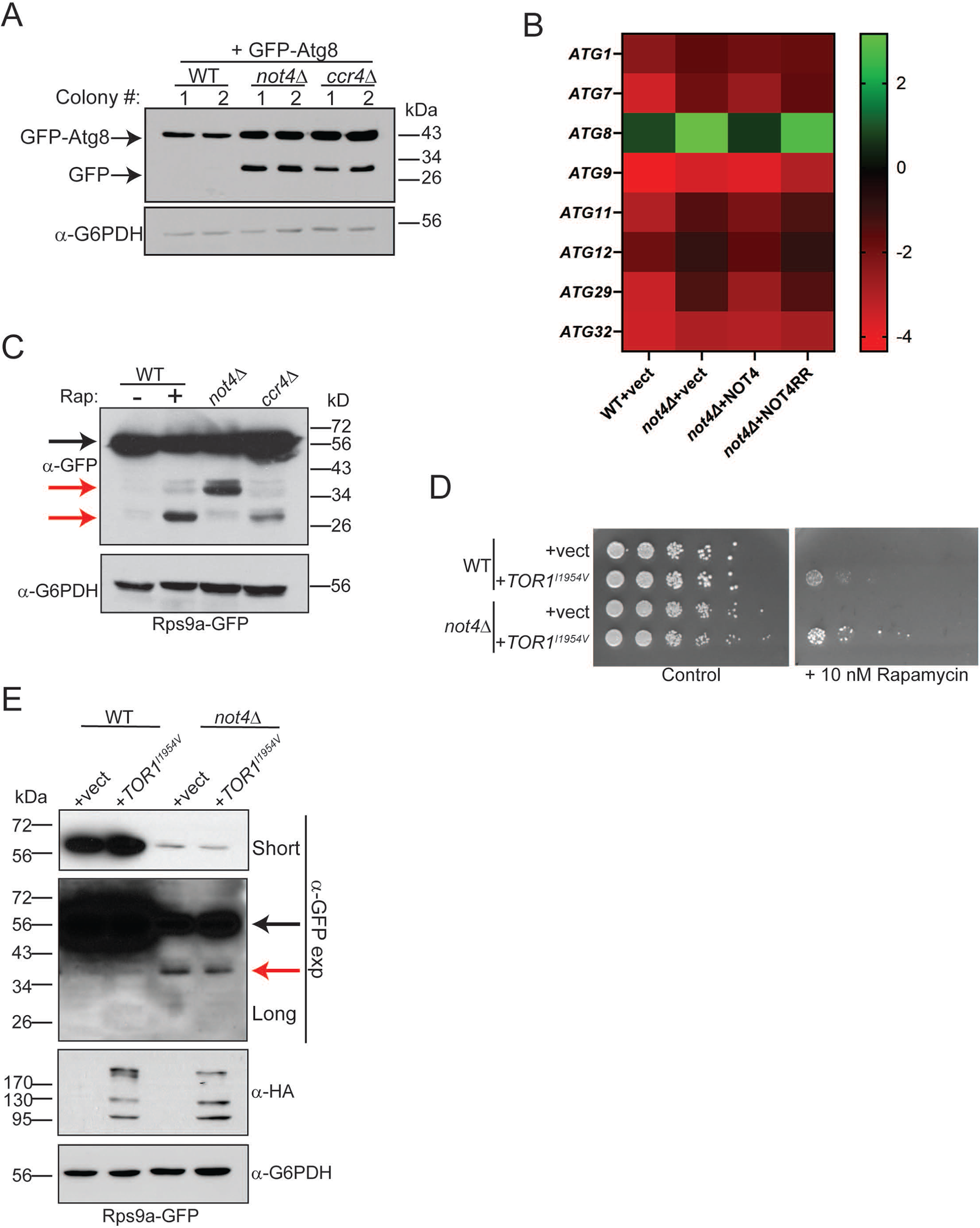
Ccr4-Not ligase deficiency causes 40S RP ribophagy independent of TORC1 inhibition. **A.** A plasmid expressing GFP-Atg8 from its native promoter was transformed into WT, *not4Δ*, and *ccr4Δ*. Log phase WCEs from two independent transformants for each strain background were analyzed by α-GFP IB. The GFP-Atg8 and GFP only bands are indicated. **B.** RT-qPCR analysis of the indicated *ATG* mRNAs was performed from three independent biological replicates normalized to the control gene *SPT15*. The average signal for all three replicates was Log_2_ converted and are presented as a heat map. Bar graphs for the mean and SD of each individual *ATG* mRNA are in Figure S7. **C.** The WT, *not4Δ*, and *ccr4Δ* strains expressing Rps9a-GFP were grown to log phase before preparing WCEs. In parallel, a log phase WT culture was treated with 200 nM rapamycin (Rap) for fours before harvesting and WCE preparation. Samples (30 µg) were analyzed by α-GFP IB and then α-G6PDH as a loading control. The black arrow indicates full-length Rps9a-GFP and the red arrows indicate partial or complete (∼ 27 kDa) degradation products. **D.** WT and *not4Δ* were transformed with control vector or vector expressing the Tor1^I1954V^ mutant. Equal numbers of cells from overnight cultures were 5-fold serially diluted and spotted to control selection plates or selection plates containing the TORC1 inhibitor rapamycin (10 nM). Images were acquired after four days of growth at 30°C. **E.** The strains from **D** were grown to log phase and WCEs (30 µg) then analyzed by α-GFP IB to detect Rps9a-GFP and α-HA to detect Tor1^I1954V^ expression. The α-G6PDH IB is the loading control. Short and long α-GFP IBs are indicated with the black arrow denoting full-length Rps9a-GFP and the red arrow indicating the partial degradation product.

The macroautophagy upregulation in *not4Δ* and *ccr4Δ* suggested the partial or complete Rps9a degradation in these cells could be due to increased 40S ribophagy since it is proteasome-independent (Figure 4). To test this possibility, Rps9a-GFP WT, *not4Δ*, and *ccr4Δ* cultures in nutrient-rich media were harvested, while a duplicate WT culture was treated with the TORC1 inhibitor rapamycin for four hours to induce ribophagy. No Rps9a-GFP degradation occurred in the WT control, while substantial free GFP (∼ 27 kDa) accumulated in both the rapamycin-treated WT and *ccr4Δ* cells (Figure 5C) . Importantly, an intermediate Rps9a-GFP degradation product was present in the WT rapamycin treated sample that matched the size of the partial degradation product in *not4Δ* (Figure 5C). These data support a role for 40S ribophagy upregulation in Ccr4-Not deficient cells. Because loss of either Ccr4 or Not4 inhibits TORC1 signaling, we tested if the 40S ribophagy in *not4Δ* was due solely to decreased TORC1 signaling. WT and *not4Δ* were transformed with a control plasmid or plasmid expressing an HA-tagged Tor1^I1954V^ mutant that increases Tor1 kinase activity. Tor1^I1954V^ rescued WT and *not4Δ* growth in the presence of rapamycin (Figure 5D), thus confirming it enhances TORC1 signaling in both cell types. However, Tor1^I1954V^ failed to restore Rps9a-GFP expression or prevent 40S ribophagy in *not4Δ* (Figure 5E). Therefore, Ccr4-Not inhibits 40S ribophagy in nutrient-rich environments through a mechanism that depends on both its ubiquitin ligase and deadenylase subunits, and this 40S ribophagy inhibition is not due solely to Ccr4-Not ubiquitin ligase-dependent TORC1 activation.

Previously, *not4Δ* was reported to have a minor synthetic sick phenotype when combined with an *atg7Δ* mutant that blocks autophagosome-dependent macroautophagy (23). Importantly, no increase in 40S ribophagy was reported in this study, although a different 40S reporter strain (Rps3-GFP) was used and a non-specific GFP-reactive band was present at a size similar to the partial Rps9a-GFP degradation product we detect in *not4Δ* (Figure 3C and Figure 5C). We revisited the possibility that the partial 40S ribophagy in *not4Δ* could be explained by increased Atg7-dependent macroautophagy (Figure 5) by combining *not4Δ* with *atg7Δ*. As previously reported, the *atg7Δ not4Δ* exhibited only a mild synthetic sick phenotype compared to *not4Δ* (Day 2 in Figure 6A). However, the *not4Δ atg7Δ* neither restored full-length Rps9a-GFP expression nor prevented Rps9a-GFP partial degradation (Figure 6B and 6C). Thus, while Ccr4-Not ligase disruption does deregulate macroautophagy in nutrient rich environments (Figure 5A), the 40S RP repression and 40S ribophagy caused by *not4Δ* are macroautophagy-independent.

**Figure 6.**
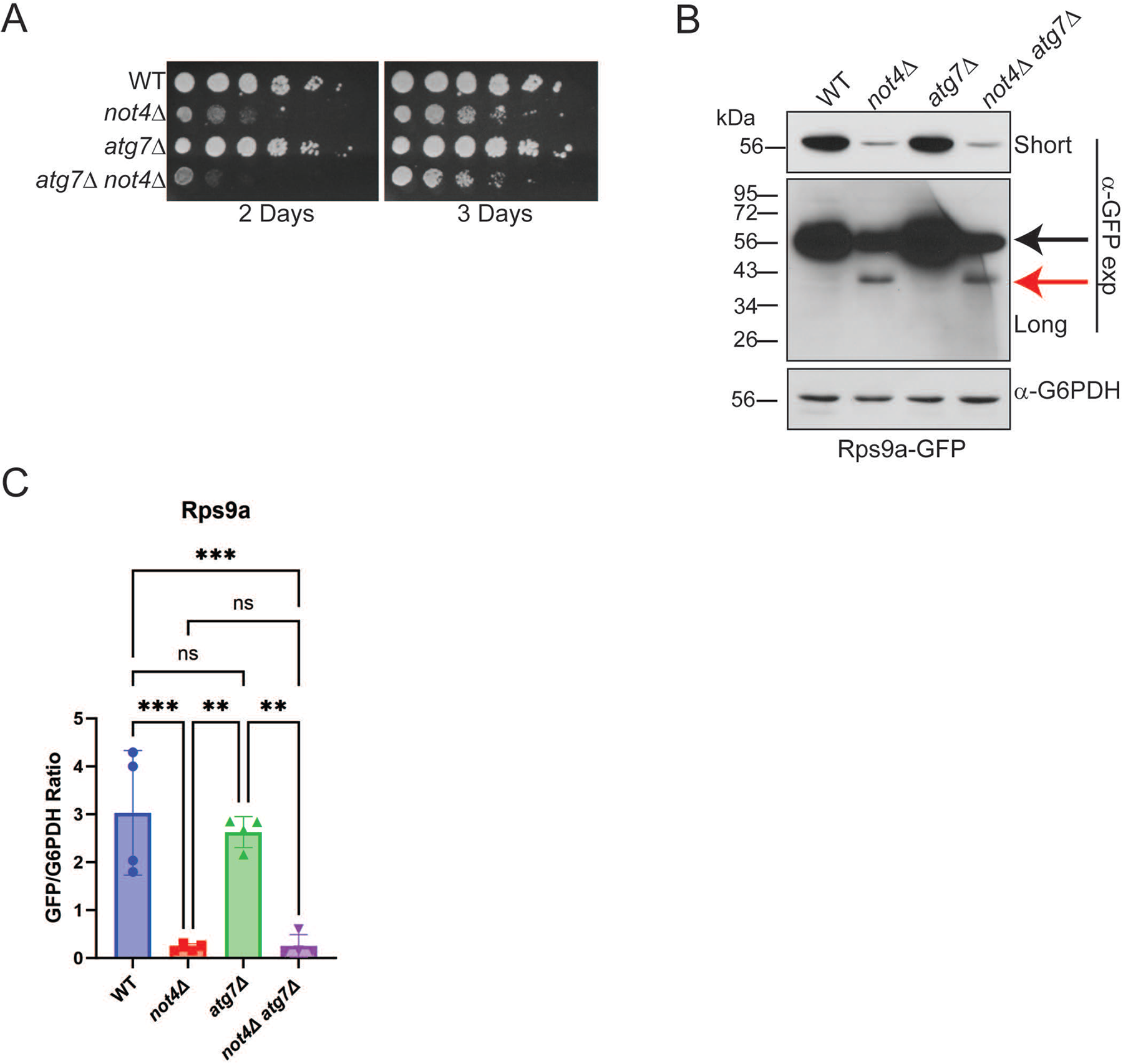
40S ribophagy activation in Ccr4-Not ligase deficient cells is independent of macroautophagy. **A.** The individual and double mutants were cultured overnight and then equal cell numbers were 5-fold serially diluted, spotted to YPD plates, and incubated at 30°C. Images were taken at the indicated times. **B.** Log phase WCEs (30 µg) from the strains in **A** were analyzed by α-GFP IB. Short and long exposures are provided with the full-length Rps9a-GFP indicated by the black arrow and the degradation product denoted by the red arrow. **C.** Quantification of **B**. The short α-GFP IBs for a total of 3-4 individual biological replicates were quantified and expressed as the ratio of full-length GFP to G6PDH. *\*\*-p< 0.005* or greater by one-way ANOVA.

### 40S ribophagy requires the ESCRT pathway in wild-type cells but additional pathways mediate 40S ribophagy in a Ccr4-Not ligase mutant

The effectors selectively regulating 40S ribophagy are unknown (52,62). Because the *not4Δ* vector and *NOT4RR* cells both induced partial 40S ribophagy, we inspected the DEPs overlapping in these cells (Figure 1D) for candidate proteins connected to autophagy or ribosome turnover for possible 40S ribophagy effectors. Three upregulated proteins were identified (Cue5, Nvj1, and Vps27) that have established roles in various autophagy pathways (Figure S8). Cue5 is a CUET protein family member that binds ubiquitin and interacts with Atg8 to mediate proteaphagy and protein aggregate autophagy (63–65). Nvj1 is a nuclear membrane protein that interacts with the vacuole-localized protein Vac8 to promote piecemeal microautophagy of the nucleus (PMN) during nutrient stress (66). Vps27 is an endosomal sorting complex required for transport (ESCRT) subunit that binds ubiquitinated substrates to promote their degradation via autophagosome-independent microautophagy (67). Vps27 also regulates endosomal protein sorting and turnover of plasma membrane proteins (68). Transcript analysis by RT-qPCR revealed that loss of Not4 ligase activity increased the mRNA levels for all three proteins (Figure 7A), indicating that Not4 ligase activity either promotes their mRNA decay or represses their transcription. To determine if they contribute to 40S ribophagy, WT, *cue5Δ*, *nvj1Δ*, and *vps27Δ* were either mock treated or treated with rapamycin for four hours to induce ribophagy before analyzing Rps9a-GFP cleavage. While no Rps9a-GFP cleavage occurred in the mock-treated cells, TORC1 inhibition induced robust Rps9a-GFP cleavage in WT, *cue5Δ*, and *nvj1Δ*, while this was prevented by *vps27Δ* (Figure 7B). These data support a role for the Vps27-dependent ESCRT pathway in nutrient-stress induced 40S ribophagy. 60S ribosomes also undergo nutrient-stress induced ribophagy. This process is regulated by the Ltn1 ubiquitin ligase that ubiquitinates the 60S to block 60S ribophagy, and the Ubp3 and Bre5 (Ubp3/Bre5) deubiquitinase complex that deubiquitinates the 60S to activate 60S ribophagy (62,69). Because 60S ribophagy was not detected in *not4Δ* or *ccr4Δ* (Figure 3C), we inspected the Not4 proteome data to determine if the expression of these 60S ribophagy effectors was altered. Intriguingly, while Ltn1 and Ubp3 protein levels were not affected (fold change less than 1.5), we found that Bre5 was repressed significantly and it is among the 422 proteins that overlap in the *not4Δ* vector and *NOT4RR* cells (Figure S9 and Figure 1D). Overall, these data demonstrate that the ubiquitin-binding ESCRT subunit Vps27 contributes to 40S ribophagy activation, suggesting the possibility that Vps27 upregulation in Ccr4-Not ligase-deficient cells may activate 40S ribophagy. Furthermore, our proteomic data also suggest a possible explanation for why 60S ribophagy is not induced in Not4-deficient cells since Bre5 expression is repressed.

**Figure 7.**
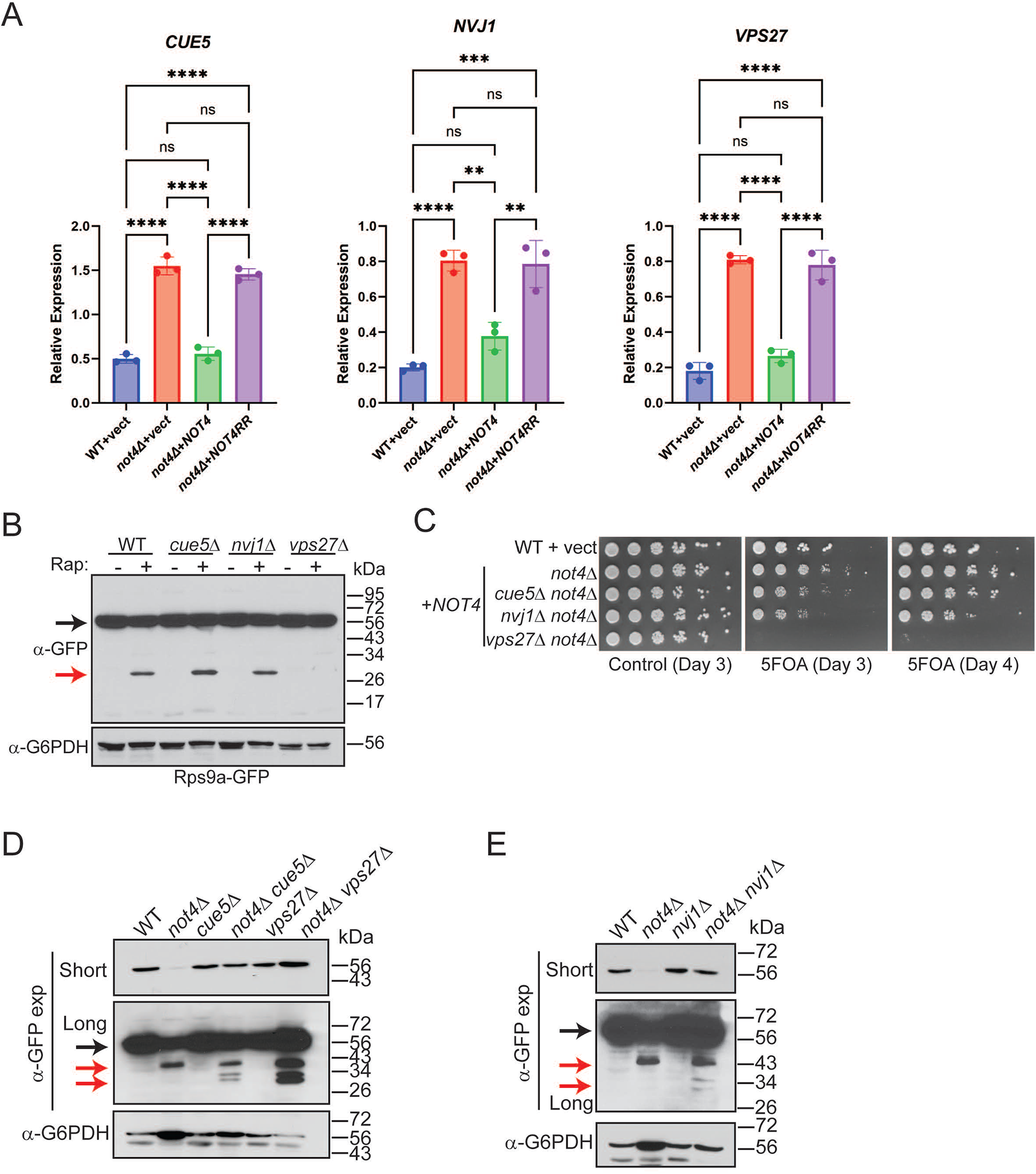
Both 40S RP repression and ribophagy activation in Ccr4-Not ligase deficient cells are functionally connected to the endolysosomal pathway. **A.** RT-qPCR analysis of the indicated genes from triplicate independent biological replicates with the mean and SD plotted and significance determined by one-way ANOVA. *\*\*-p<0.005* or greater. **B.** The WT and individual mutants were cultured to log phase and mock treated or treated with 200 nM rapamycin for four hours before harvesting and performing α-GFP IB. Black arrow indicates full-length Rps9a-GFP and the red arrow indicates free GFP. **C.** WT cells and *not4Δ* transformed with either control or *NOT4* expression vector were cultured overnight. Equal cell numbers then were 5-fold serially diluted, spotted to plasmid-selective control media or to media contain 5-FOA, and then incubated at 30°C for the indicated times before photographing. **D** and **E**. The WT, single, and double mutants were grown to log phase and then analyzed by α-GFP IB. Short and long α-GFP exposures are provided. The black arrow indicates full-length Rps9a-GFP and the red arrows indicate partial and complete GFP degradation products.

If Not4-deficient cells increase Vps27 expression as the sole mechanism to induce 40S ribophagy, then disrupting this pathway in *not4Δ* should prevent 40S ribophagy activation. To test this possibility, *not4Δ* was combined separately into *vps27Δ*, and also into *cue5Δ* or *nvj1Δ*, and then the growth phenotypes of the single and double mutants were compared. The *cue5Δ not4Δ* and *nvj1Δ not4Δ* exhibited synthetic sick phenotypes when forced to lose the *NOT4* expressing plasmid on 5-FOA plates, while the *vps27Δ not4Δ* was profoundly more growth impaired (Figure 7C). This extreme synthetic sick phenotype is consistent with a previous report that found mating *vps27Δ* to *not4Δ* failed to generate viable *vps27Δ not4Δ* spores. Surprisingly, when Rps9a-GFP expression was examined in the single and double mutants, each double mutant restored Rps9a-GFP levels back to WT (Figure 7D and 7E). Furthermore, none of the double mutants, including *vps27Δ not4Δ*, blocked Rps9a-GFP ribophagy. Instead, each double mutant increased conversion of the partial Rps9a-GFP degradation product in *not4Δ* to free GFP (Figure 7D and 7E), indicating that loss of these autophagy factors partially alleviates the block in 40S ribophagy caused by *not4Δ*. Therefore, although WT cells require Vps27 for 40S ribophagy during nutrient stress, redundant mechanisms must activate 40S ribophagy in *not4Δ* .

## DISCUSSION

Cells coordinate anabolism and proliferation with their environment, which involves communication between the endolysosomal pathway and the machinery controlling gene expression, translation, and proteostasis. The Ccr4-Not complex regulates each of these processes, so it is uniquely positioned to integrate communication between these interrelated activities. Ccr4-Not’s role in transcriptome regulation is known, yet how Ccr4-Not ubiquitin signaling affects the proteome is incompletely understood. We have addressed this issue by performing proteomic analysis of cells lacking Not4, expressing WT Not4, or expressing a full-length inactive Not4 mutant. This approach controls for proteome alterations indirectly caused by Ccr4-Not structural changes that could affect its non-ligase signaling roles when Not4 is absent. Indeed, *not4Δ* vector and *not4Δ NOT4RR* cells share many DEPs (422 in total) that likely represent proteins regulated by Not4 ligase activity. However, *not4Δ* vector cells also have approximately 300 DEPs that are not explained by loss of Not4 ligase function. These differences may reflect indirect effects on non-ligase dependent Ccr4-Not regulated activities, or they may indicate that Not4 contributes to mRNA transcription, stability, and/or mRNA translation independently of its ubiquitin ligase role. We also find that increased Not4 levels upregulate mitochondrial proteins, indicating that Not4 ligase signaling may mediate mitochondrial metabolic reprogramming, although how this occurs is not yet understood. There is previous support for a Ccr4-Not mediated role in multiple aspects of mitochondrial regulation (29,41,42,70), so future studies will need to address these important questions.

The two most intriguing findings presented herein are the RP and Rib repression due to dysregulated Ccr4-Not ligase activity, and the activation of 40S ribophagy when Ccr4-Not dependent ubiquitin signaling is lost. The repressed GOs for each experimental condition were dominated by ribosomal categories, clearly indicating Not4 ligase signaling is required for ribosome homeostasis. Not4 loss disrupts polysome formation and alters ratios of 40S, 60S, and 80S ribosome. Additionally, a previous metabolomic analysis found that the *not4Δ* metabolic profile resembled that of ribosomal mutants (71). These previous studies, and our new data, thus provide increased support for the concept that Not4 maintains ribosomal homeostasis. While Not4 overexpression restores many RP and Ribi factors, a core set remain repressed and overlap those inhibited in *not4Δ* vector and *NOT4RR* cells (see Figure S6 for visual representation). However, since Not4 overexpression restores the protein synthesis defects and sensitivity to translational stress in *not4Δ*, their repression is insufficient to inhibit global protein synthesis. Another important observation is that the RP repression we detect is not selective for either 40S or 60S RPs but instead it encompasses both. Whether the mechanisms repressing both 40S and 60S RPs are the same or differ will need to be addressed in future studies, but it is clear this repression occurs post-transcriptionally. Our analysis of the model RP Rps9a indicates that neither proteasome nor macroautophagy inhibition restores Rps9a expression in *not4Δ*. Therefore, Not4 ligase activity likely does not inhibit RPs by enhancing their degradation through these pathways.

We believe the most likely explanation for the observed Not4 RP and Ribi inhibitory effects is that Ccr4-Not ligase signaling controls RP and Ribi mRNA translation. Ccr4-Not can bind to RP mRNAs during their transcription elongation, which affects their cytoplasmic translation (72). If this process is dependent on Not4 ligase activity, then altering Not4 dosage or function may impair RP mRNA translation. A Not4 dosage and ligase-dependent inhibitory effect on RP expression is supported by our targeted analysis of Rps9a and Rpl36a since Not4 expressed from its native regulatory elements restores their normal expression. Increased Not4 dosage in WT cells also enhances sensitivity to selective translational stressors that impair ribosome activity at the A-site site. These genetic data may indicate that factors interacting with the ribosome at or near the A-site could be sensitive to Not4 ligase signaling. In total, these data indicate support for the concept that balanced Not4 ligase signaling maintains ribosomal homeostasis.

Another key finding is that either Not4 or Ccr4 loss activates 40S ribophagy, indicating they likely repress 40S ribophagy through a shared mechanism requiring Ccr4-Not ubiquitin signaling. Future efforts will need to address if Ccr4 inhibits 40S ribophagy in a deadenylase-dependent or independent manner. However, Ccr4 does regulate endolysosomal nutrient signaling independently of its deadenylase activity (29), and Ccr4 also participates with Not4 in some aspects of proteome regulation (73). Therefore, precedents exist for Ccr4 to have non-mRNA degradative activities. In addition, the deregulated macroautophagy in Ccr4-Not mutants does not explain the 40S ribophagy since *atg7Δ not4Δ* fails to prevent it. It also is not due solely to inhibition of TORC1 since restoring TORC1 signaling does not prevent 40S ribophagy or restore RP expression in *not4Δ*. Interestingly, whereas *not4Δ* allows only partial 40S RP degradation, free GFP can be liberated in *ccr4Δ*. Previously, we found that Ccr4-Not activates TORC1 by promoting vacuole V-ATPase activity, and *ccr4Δ* reduces vacuole acidification (29). Our unpublished data indicate that *not4Δ* inhibits V-ATPase dependent nutrient signaling more than *ccr4Δ.* Therefore, the incomplete 40S ribophagy in *not4Δ* likely is due to greater V-ATPase inhibition that increases vacuole pH even more than *ccr4Δ* and inhibits the proteolytic activities required for ribosome degradation. This possibility is supported by our genetic data demonstrating that loss of the plasma membrane pump Pdr5, or the vacuole Na^+^-K^+^/H^+^ exchanger Nhx1, alleviates both the 40S ribophagy block and restores RP expression. These mutants likely compensate for the impaired V-ATPase in *not4Δ* by reducing competition for limiting H^+^ or by decreasing vacuole H^+^ efflux. Such an effect would enhance vacuole protein degradation and activation of the downstream nutrient signaling controlling RP expression.

The independence of *not4Δ* 40S ribophagy from macroautophagy indicates that the mechanistically distinct microautophagy pathway is responsible. K48 and K63 linked substrate polyubiquitination can signal degradation through both the macroautophagy and microautophagy pathways (21,22). Our data indicates that K48-linked ubiquitin chains accumulate on Rps9a in *not4Δ*, thus suggesting this ubiquitination may flag the 40S for ribophagy similar to that which occurs for proteaphagy. How K48-linked 40S ubiquitination accumulates is not known, but it may be linked to the defects in proteasome-dependent deubiquitination in Not4 deficient cells. Ribosome downregulation can mediate adaptation to proteostatic stress (74), so the combined RP/Ribi repression and 40S ribophagy activation in Ccr4-Not mutants may allow them to adapt to the increased proteostatic stress caused by proteosome dysregulation.

Three autophagy effectors (Cue5, Nvj1, and Vps27) are upregulated in *not4Δ* vector and *NOT4RR* cells where 40S ribophagy is activated. Importantly, only loss of Vps27, which is an ESCRT ubiquitin binding subunit controlling microautophagy, blocked nutrient stress-induced 40S ribophagy in wild-type cells. To our knowledge this is among the first 40S ribophagy effectors identified. However, 40S ribophagy was not prevented by a *vps27Δ not4Δ* mutant, thus indicating additional, and likely redundant, pathways mediate 40S ribophagy activation in Ccr4-Not mutants. Surprisingly, the 40S RP degradation defect in *not4Δ* was partially restored by *vps27Δ not4Δ*, and also by *cue5Δ not4Δ* and *nvj1Δ not4Δ*. How co-disruption of these pathways enhances 40S RP degradation is not understood, but their effect appears similar to that which occurs upon loss of Pdr5 or Nhx1. A common theme connecting them is their linkage to the endolysosomal pathway. Cue5 is best defined as a ubiquitin-binding receptor for proteaphagy and aggrephagy (63,64), yet it has extensive genetic interactions with endolysosomal factors that suggests it has endolysosomal functions not yet identified (39,75). Cue5 loss in *not4Δ* may affect these other activities to restore RP expression and enhance 40S ribophagy. Although Nvj1 mediates PMN, it also interacts with the oxysterol binding protein Osh1 that promotes nutrient transport from the plasma membrane (66). A *nvj1Δ* can enhance nutrient uptake by not sequestering Osh1 from this additional role (76). The *vps27Δ* may increase nutrient transporters, and possibly H^+^ exporters, on the cell surface since their turnover would be reduced through the endolysosomal pathway (17). Thus, the vacuole defects in *not4Δ* may be partially compensated by loss of these additional autophagy related factors.

During the course of this study, a SILAC-based analysis of the WT and *not4Δ* proteome found that *not4Δ* increases expression of several RPs relative to WT cells (31), which is a result seemingly at odds with our proteomic data. Previous studies using specialized protein extraction approaches found that *not4Δ* can form protein aggregates *in vivo*, which may be related to loss of Rps7a monoubiquitination (23,24). However, to prevent aggregates we prepared the proteomic samples by bead beading in high salt buffer. We also directly demonstrate that both Rps9a and Rpl36a are repressed in *not4Δ* even when analyzed under denaturing conditions. Depletion of aggregated proteins from the extracts used in our proteome analysis also would not account for the significant RP and Ribi repression in *NOT4* expressing cells where Rps7a monoubiquitination is intact, and it also would not explain why disrupting endolysosomal effectors in *not4Δ* restores RP expression. Future efforts will be required to reconcile the differences between our results and this recent report.

Analysis of Ccr4-Not has focused predominantly on its role in transcription and mRNA decay. However, increasing evidence indicates that its ubiquitin signaling role is critically important to additional cellular processes. The data presented herein demonstrate that balanced Not4 ligase activity is required to maintain RP expression, while Ccr4-Not mediated ligase signaling also inhibits 40S ribophagy. Combined with our previous results connecting Ccr4-Not to endolysosomal regulation, these new data reveal an important role for the Ccr4-Not ligase in endolysomal-dependent nutrient signaling and autophagy control than currently is appreciated. Since Ccr4-Not dysregulation is implicated in many diseases and developmental disorders, probing these additional Ccr4-Not regulated processes will be essential to understanding Ccr4-Not’s role in these conditions.

## DATA AVAILABILITY

The mass spectrometry proteomics data have been deposited to the ProteomeXchange Consortium via the PRIDE partner repository with the dataset identifier PXD044420 and 10.6019/PXD044420 (Reviewer account details: **Username:**, **Password:** 4tCDgHXm) (77,78).

## ACKNOWLEDGEMENTS

We would like to thank Drs. Daniel Klionsky and Ted Powers for their kind gifts of plasmids.

