## Supplemental Figures for "Ccr4-Not ubiquitin ligase signaling regulates ribosomal protein homeostasis and inhibits 40S ribosomal autophagy"

### SUPPLEMENTAL FIGURE LEGENDS

**Supplemental Figure 1. DEP pathway enrichment analysis of WT + vector control relative to *not4Δ* + vector expressing cells.** Dot blot analysis of the cellular component GO categories overrepresented in the down regulated proteins (**A**) and up regulated proteins (**B**).

**Supplemental Figure 2. DEP pathway enrichment analysis of WT + vector control relative to *not4Δ* + *NOT4* expressing cells.** Dot blot analysis of the cellular component GO categories overrepresented in the down regulated proteins (**A**) and up regulated proteins (**B**).

**Supplemental Figure 3. DEP pathway enrichment analysis of WT + vector control relative to *not4Δ* + *NOT4RR* expressing cells.** Dot blot analysis of the cellular component GO categories overrepresented in the down regulated proteins (**A**) and up regulated proteins (**B**).

**Supplemental Figure 4. RT-qPCR analysis of RP genes. A** and **B** are the mean and SD of three independent biological replicates with the RP gene-specific signal normalized to the internal reference control gene *SPT15*. The mean for each of these genes was $Log_2$ transformed and used to generate the heat maps in **Figure 2**.

**Supplemental Figure 5. Proteomic quantification of the known Not4 substrates Rps7a, Egd1, and Egd1.** The mean and SD of the normalized protein abundance for each indicated protein is plotted. The fold change relative to the WT + vector control was less than 1.5 for each protein, and the significance of the individual replicates between each condition was analyzed by one way ANOVA. *-*$p<0.05$*.

**Supplemental Figure 6. STRING analysis of the individual Venn components.** The proteins in each component of the Venn from **Figure 1** were analyzed via the STRING database using a 0.9 confidence setting with the unconnected nodes removed. The arrows align the remaining connected networks to their respective Venn section. Only those networks related to RP and Ribi function are labeled.

**Supplemental Figure 7. RT-qPCR analysis of *ATG* genes. A** and **B** are the mean and SD of three independent biological replicates with the *ATG* gene-specific signal normalized to the internal reference control gene *SPT15*. The mean for each of these genes was $Log_2$ transformed and used to generate the heat maps in **Figure 5**.

**Supplemental Figure 8. Proteomic quantification of the autophagy factors Cue5, Nvj1, and Vps27.** The mean and SD of the normalized protein abundance for each indicated protein is plotted. The fold increase relative to the WT + vector control for Cue5, Nvj1, and Vps27 in the *not4Δ* + vector and *not4Δ* + *NOT4RR* samples was 1.5 fold or higher. The significance of the individual replicates between each condition was analyzed by one way ANOVA. *-*$p<0.05$*.

**Supplemental Figure 9. Proteomic quantification of the 60S ribophagy factors Ltn1, Ubp3, and Bre5.** The mean and SD of the normalized protein abundance for each indicated protein is plotted. The fold change relative to the WT + vector control for Ltn1 and Ubp3 was less than 1.5 fold in all the experimental conditions. Bre5 expression was reduced in the *not4Δ* + vector (-1.96 fold) and it was reduced in the *not4Δ* + *NOT4RR* (-1.75 fold). The significance for the individual replicates between each condition was analyzed by one way ANOVA. *-$p<0.05$.

A

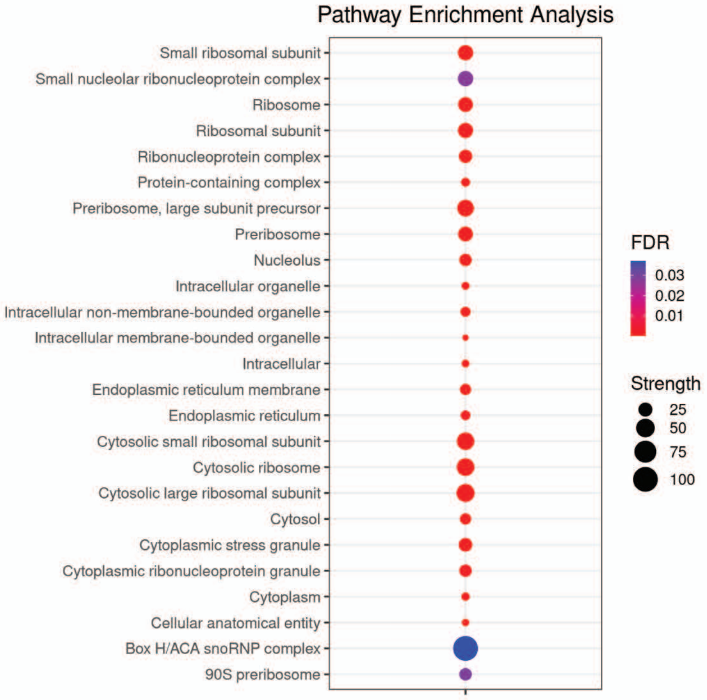


B

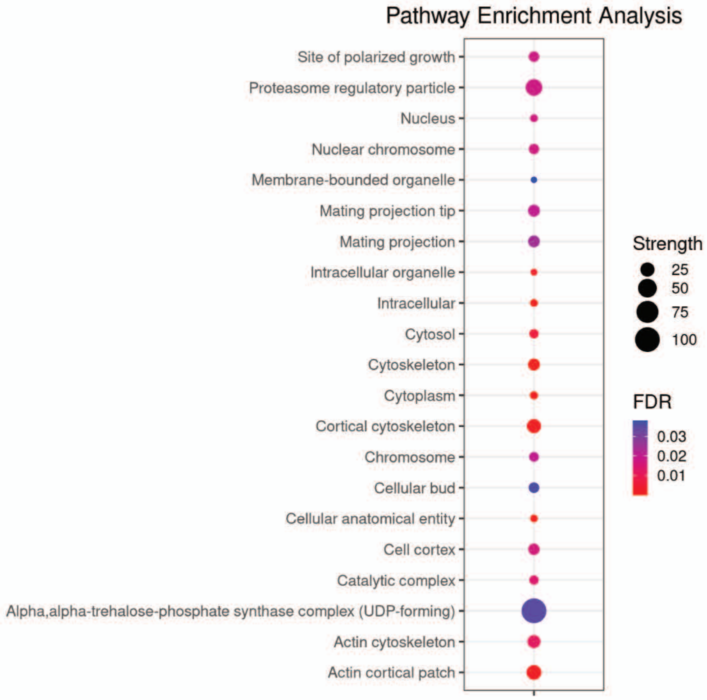


Supplemental Figure 1

A

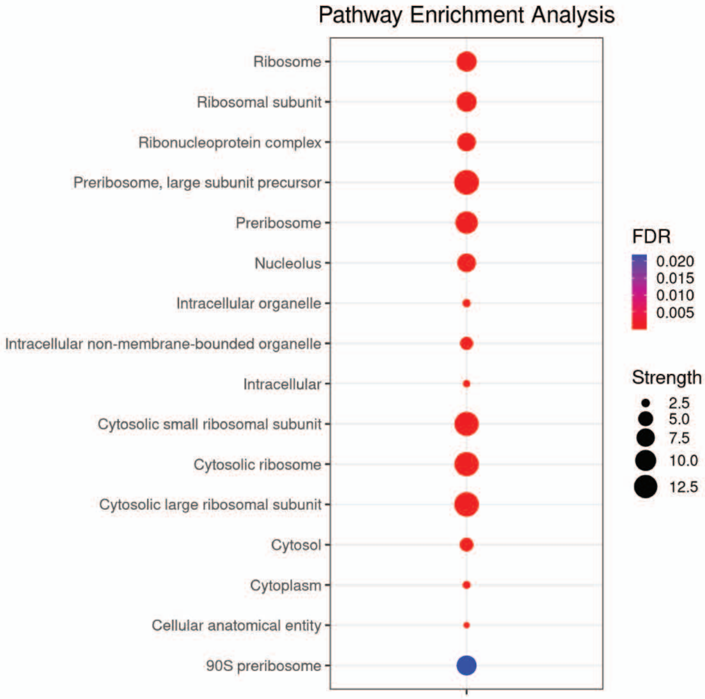


B

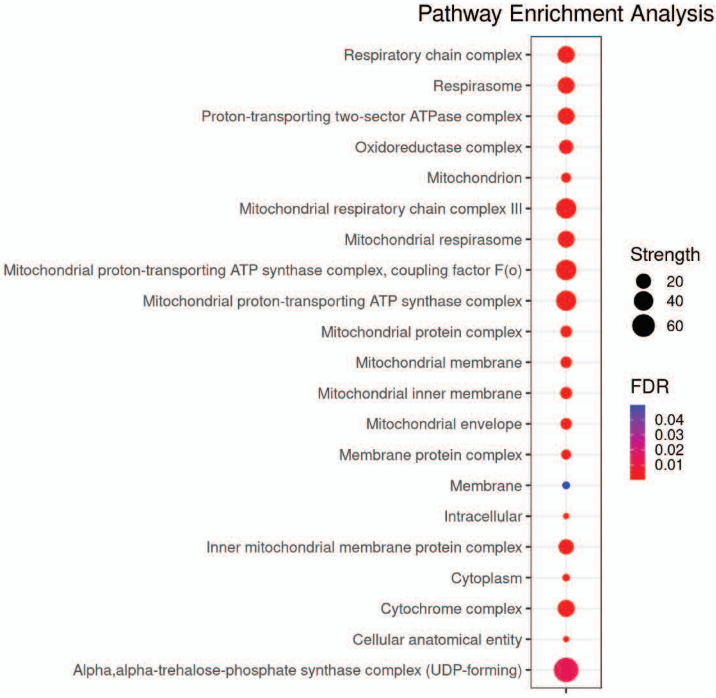


Supplemental Figure 2

A

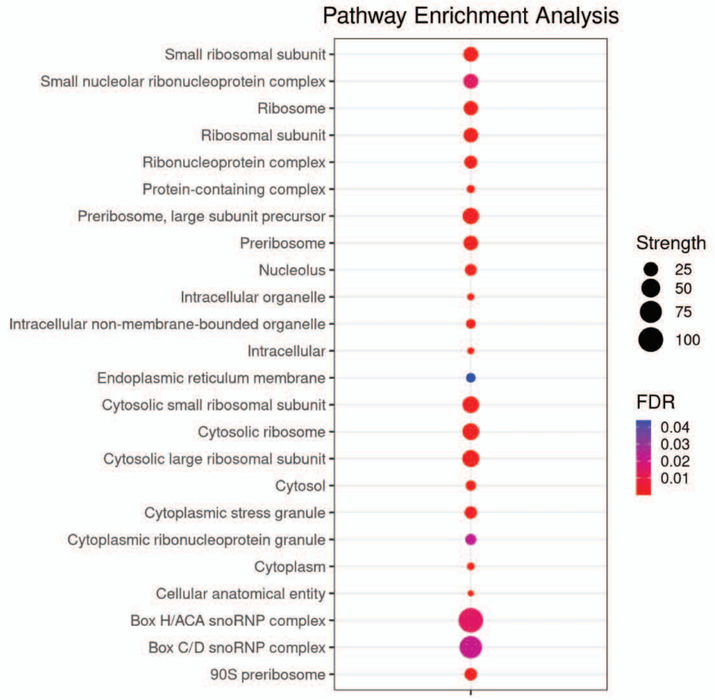


B

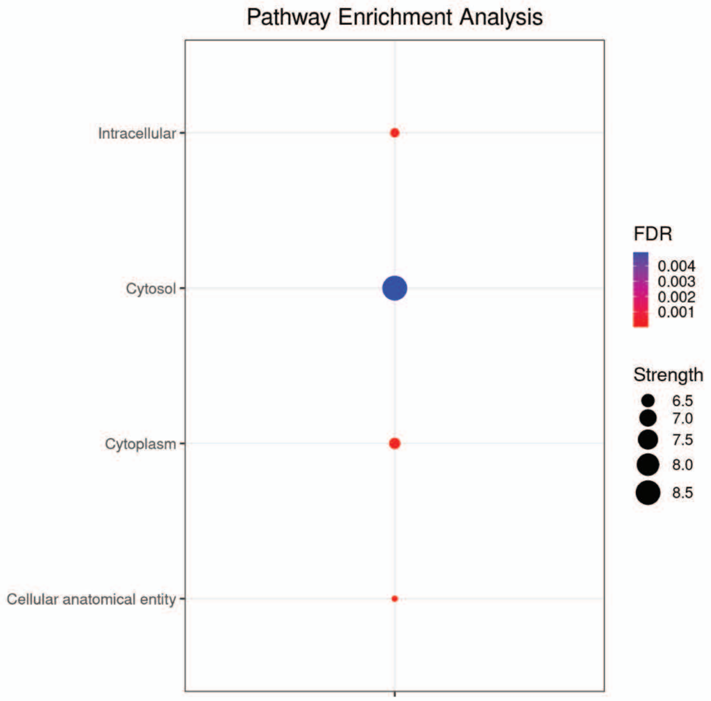


Supplemental Figure 3

A

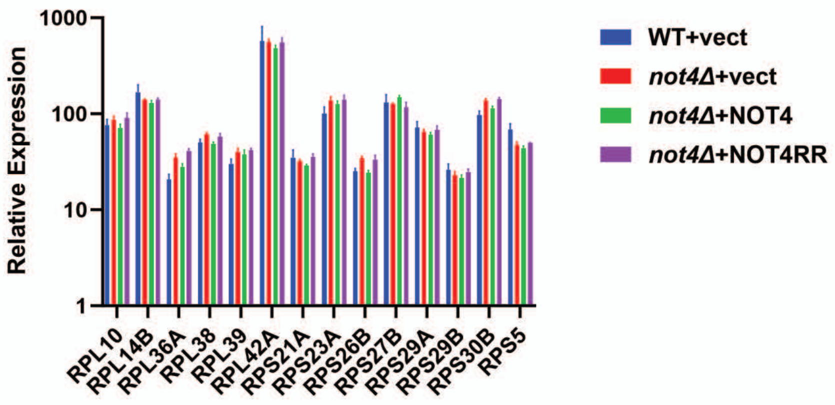


B

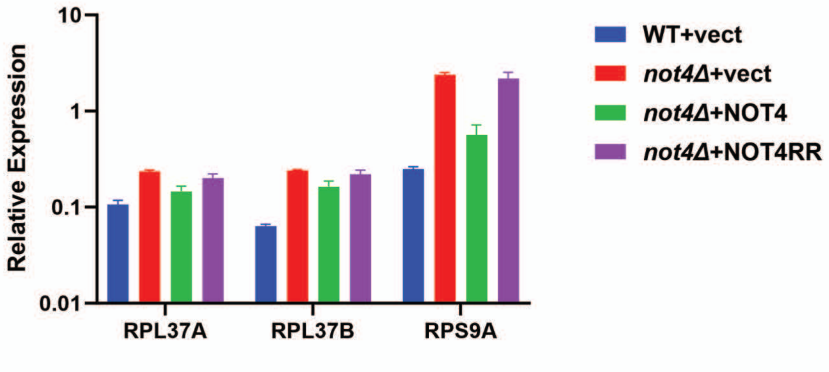


Supplemental Figure 4

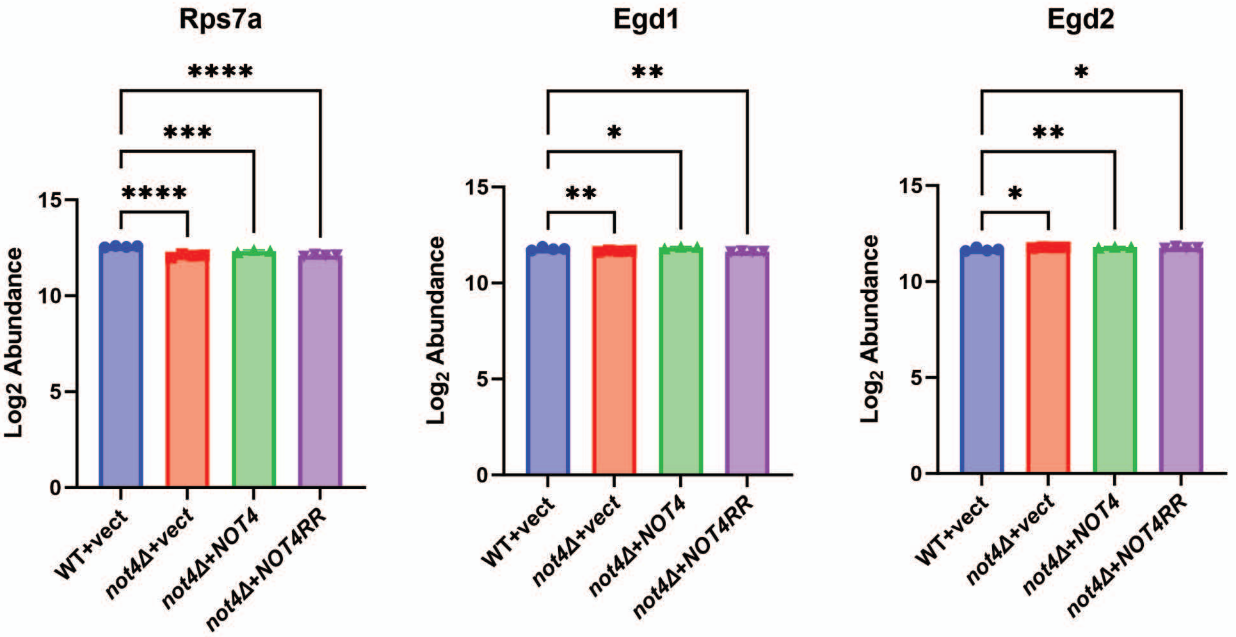


Supplemental Figure 5

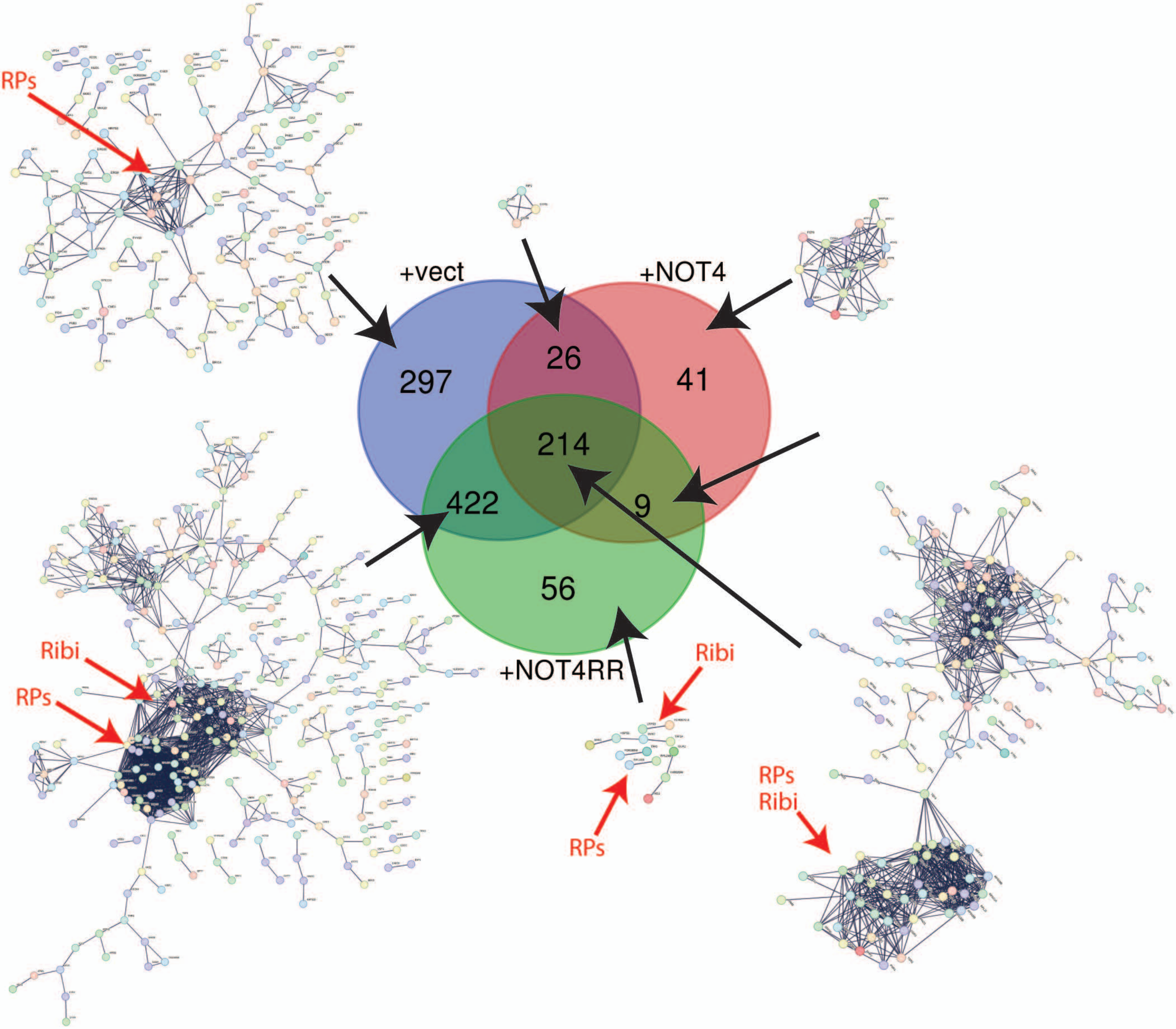


Supplemental Figure 6

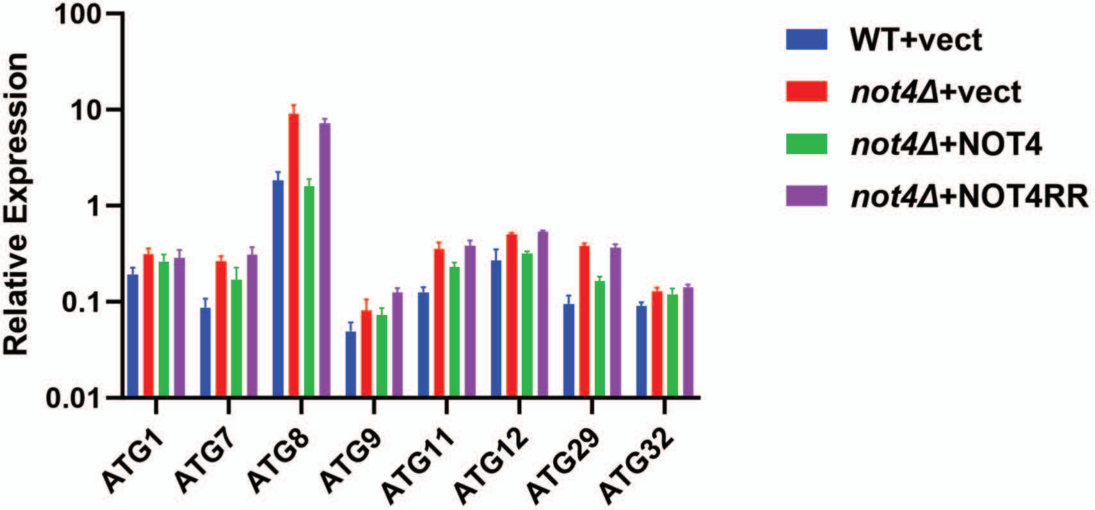


Supplemental Figure 7

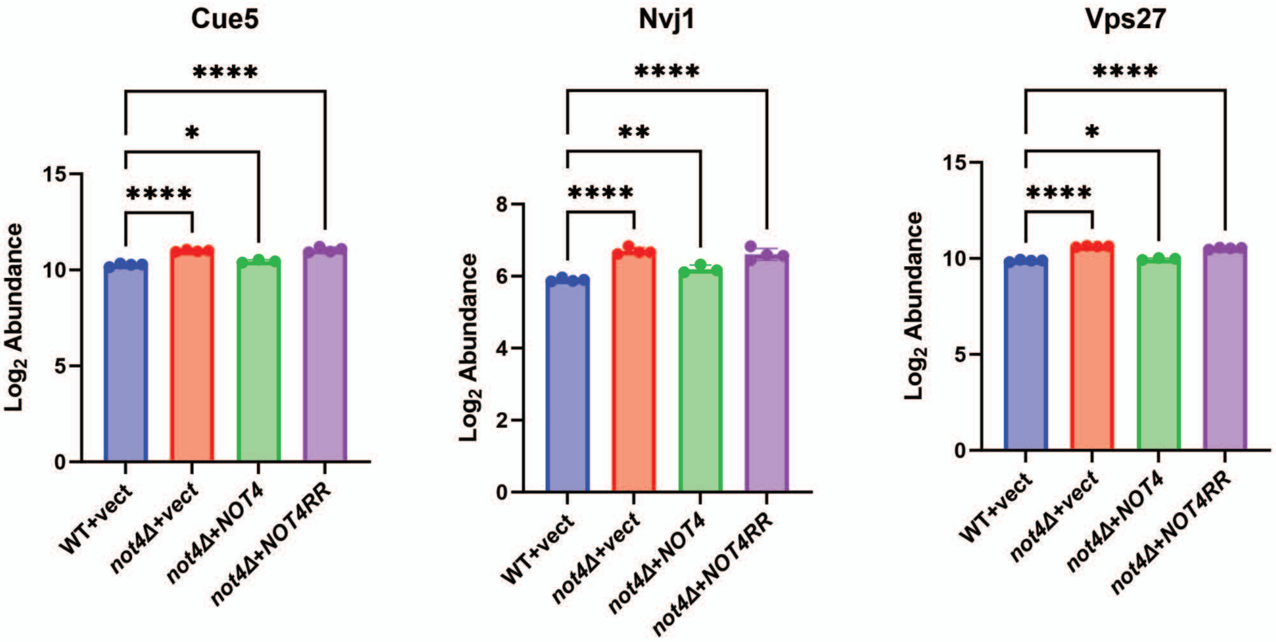


Supplemental Figure 8

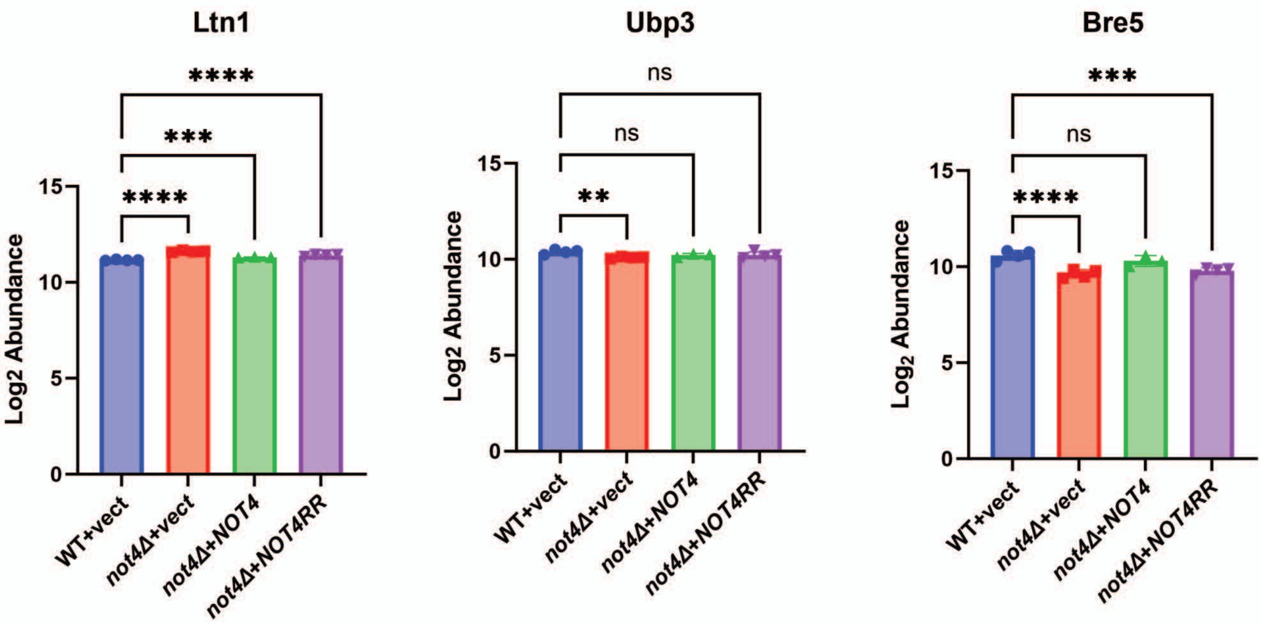


Supplemental Figure 9
