## Supplementary material for "Ccr4-Not ubiquitin ligase signaling regulates ribosomal protein homeostasis and inhibits 40S ribosomal autophagy": Table S1

**Supplemental Table 1. Yeast strains.**

| **Strain** | **Description** | **Reference** |
| --- | --- | --- |
| BY4741 | *MATa his3-D1 leu2-D0 met15-D0 ura3-D0* | Dharmacon/  Open Biosystems |
| *ccr4Δ* | *MATa his3-D1 leu2-D0 met15-D0 ura3-D0 ccr4D::K KanMX* | Dharmacon/  Open Biosystems |
| *not4Δ* | *MATa his3-D1 leu2-D0 met15-D0 ura3-D0 not4D::K KanMX* | Dharmacon/  Open Biosystems |
| YNL992 | *MATa his3-D1 leu2-D0 met15-D0 ura3-D0 RPS9A-EGFP::KanMX* | This study |
| YNL994 | *MATa his3-D1 leu2-D0 met15-D0 ura3-D0 RPS9A-EGFP::KanMX not4D::natNT2* | This study |
| YNL995 | *MATa his3-D1 leu2-D0 met15-D0 ura3-D0 RPL36A-EGFP::KanMX* | This study |
| YNL997 | *MATa his3-D1 leu2-D0 met15-D0 ura3-D0 RPL36A-EGFP::KanMX not4D::natNT2* | This study |
| YNL998 | *MATa his3-D1 leu2-D0 met15-D0 ura3-D0 RPS9A-EGFP::KanMX ccr4D::natNT2* | This study |
| YNL1000 | *MATa his3-D1 leu2-D0 met15-D0 ura3-D0 RPS9A-EGFP::KanMX ccr4D::natNT2* | This study |
| YNL1001 | *MATa his3-D1 leu2-D0 met15-D0 ura3-D0 RPS9A-EGFP::KanMX pdr5D::* *HphNT1* | This study |
| YNL1003 | *MATa his3-D1 leu2-D0 met15-D0 ura3-D0 RPS9A-EGFP::KanMX atg7D::* *HphNT1* | This study |
| YNL1009 | *MATa his3-D1 leu2-D0 met15-D0 ura3-D0 RPS9A-EGFP::KanMX atg7D::* *HphNT1 not4D::natNT2* | This study |
| YNL1005 | *MATa his3-D1 leu2-D0 met15-D0 ura3-D0 RPS9A-EGFP::KanMX pdr5D::* *HphNT1 not4D::natNT2* | This study |
| YNL1014 | *MATa his3-D1 leu2-D0 met15-D0 ura3-D0 RPS9A-EGFP::KanMX cue5D::* *HphNT1* | This study |
| YNL1018 | *MATa his3-D1 leu2-D0 met15-D0 ura3-D0 RPS9A-EGFP::KanMX nhx1D::HphNT1* | This study |
| YNL1020 | *MATa his3-D1 leu2-D0 met15-D0 ura3-D0 RPS9A-EGFP::KanMX nvj1D::* *HphNT1* | This study |
| YNL1022 | *MATa his3-D1 leu2-D0 met15-D0 ura3-D0 RPS9A-EGFP::KanMX vps27D::* *HphNT1* | This study |
| YNL1024 | *MATa his3-D1 leu2-D0 met15-D0 ura3-D0 RPS9A-EGFP::KanMX cue5D::* *HphNT1 not4D::natNT2* | This study |
| YNL1025 | *MATa his3-D1 leu2-D0 met15-D0 ura3-D0 RPS9A-EGFP::KanMX nhx1D::HphNT1 not4D::natNT2* | This study |
| YNL1027 | *MATa his3-D1 leu2-D0 met15-D0 ura3-D0 RPS9A-EGFP::KanMX nvj1D::* *HphNT1 not4D::natNT2* | This study |
| YNL1029 | *MATa his3-D1 leu2-D0 met15-D0 ura3-D0 RPS9A-EGFP::KanMX vps27D::* *HphNT1 not4D::natNT2* | This study |
