## Supplementary material for "Ccr4-Not ubiquitin ligase signaling regulates ribosomal protein homeostasis and inhibits 40S ribosomal autophagy": Table S2

**Table S2. Yeast Plasmids.**

| **Plasmid** | **Description** | **Reference** |
| --- | --- | --- |
| p416ADH | *AmpR CEN6/ARSH4 URA3 ADH1prom; CYC1term* | (1) |
| pRS416 | *AmpR CEN6/ARSH4 URA3* | (2) |
| pRS415 | *AmpR CEN6/ARSH4 LEU2* | (2) |
| pNOT4 | *AmpR CEN6/ARSH4 URA3 ADH1prom-NOT4-FLAG; CYC1term* | (3) |
| pNOT4RR | *AmpR CEN6/ARSH4 URA3 ADH1 prom-NOT4 I64A/G167A//F202A/C244A-FLAG; CYC1 term* | (3) |
| pNOT4ORF | *pR416; NOT4ORF (300 bp of promoter and 100 bp downstream of translational stop)* | This study |
| pPL156 | pPL132, *HA3-TOR1* I1954V | (4) |
| pRS416 GFP-ATG8 | *pRS416; ATG8prom-GFP-ATG8* | (5) |
| pPL156 | *LEU2, CEN/ARS HA3-TOR1^I1954V^* | (4) |
